# Large pan-cancer cell screen coupled to (phospho-)proteomics underscores high-dose vitamin C as a potent anti-cancer agent

**DOI:** 10.1101/2023.12.19.572293

**Authors:** Andrea Vallés-Martí, Franziska Böttger, Elysia Yau, Khadija Tejjani, Loes Meijs, Sugandhi Sharma, Madiha Mumtaz, Tessa Y. S. Le Large, Ayse Erozenci, Daniëlle Dekker, Tim Schelfhorst, Jan Paul Medema, Irene V Bijnsdorp, Jaco C Knol, Sander R Piersma, Thang V. Pham, Elisa Giovannetti, Connie R Jiménez

## Abstract

Increasing preclinical and clinical evidence has positioned high-dose vitamin C as a promising anti-cancer treatment that merits more clinical attention. Multiple cytotoxicity mechanisms have been described, including pro-oxidant effects. To contribute to the preclinical understanding of the broad pan-cancer effects of high-dose vitamin C in a global manner, we determined the IC50s of a large panel of cancer cell lines (n=51) representing 7 solid tumour types, and generated proteome data. The majority of cell lines were highly sensitive (IC50 range 0.036-10 mM, mean 1.7±0.4 mM), well below a clinically achievable dose. The proteome data (>5000 proteins per sample) showed that high intrinsic sensitivity is associated with proliferation, as indicated by functional enrichment of cell cycle and chromosomal region related terms, while lower sensitivity is linked to extracellular space, vesicle and mitochondria related biology. Moreover, (phospho-)proteome analysis of four pancreatic ductal adenocarcinoma cell lines treated with the respective IC50 concentrations of vitamin C (0.1-2 mM) revealed down-modulation of AKT-MTOR and immune suppressive signalling, while IFN-α response was enhanced upon vitamin C challenge on proteome level. Phosphoproteome data confirmed inhibition of MTOR and MAPK activities under vitamin C pressure. Altogether, our comprehensive pharmacological and (phospho-)proteome analysis is the first to assess cancer vulnerabilities and effects of vitamin C on a large cancer cell line panel and underscores the potential of high-dose vitamin C as an anti-cancer agent.

## 1. INTRODUCTION

Vitamin C (VitC) is a natural compound that has pro-oxidant and anticancer properties when administered intravenously in high doses (Böttger et al., 2021; Ngo et al., 2019; Reczek and Chandel, 2015; Schoenfeld et al., 2017; van der Reest and Gottlieb, 2016). The pro-oxidant properties are mainly due to the autoxidation of VitC, which forms hydrogen peroxide (H_2_O_2_) and reactive oxygen species (ROS) (Du et al., 2012; Michels and Frei, 2013; Schoenfeld et al., 2017). High-dose VitC can specifically target cancer cells due to their unique vulnerabilities compared to normal cells. Cancer cells have higher levels of labile iron, which can increase the formation of hydroxyl radicals (Böttger et al., 2021; Hayes et al., 2020). They often have defective mitochondria and a higher metabolic rate resulting in endogenously elevated levels of oxidative stress (Sullivan and Chandel, 2014). VitC can further increase these oxidative stress levels and thereby promote cancer cell death and inhibit metastatic dissemination (Porporato et al., 2017). Moreover, the activity of catalase, an enzyme that breaks down H_2_O_2_, is often reduced in cancer cells, allowing H_2_O_2_ to accumulate and form hydroxyl radicals (Böttger et al., 2021; Doskey et al., 2016; Ngo et al., 2019). Healthy cells, on the other hand, are equipped with molecular features to reduce oxidative stress induced by high-dose VitC.

Upon cell uptake, and in addition to its pro-oxidant and reducing capabilities, VitC can induce oxidative stress damaged DNA or reverse epithelial-to-mesenchymal transition (Ma et al., 2014; Zeng et al., 2019; Zhao et al., 2015). Other research suggests that VitC increases oxidative stress by inducing overexpression of tumour protein P53 in cancer cells (Böttger et al., 2021). Moreover, VitC has been found to inhibit the ROS-dependent MAPK/ERK signalling pathway (Su et al., 2021, 2019), which is crucial for cell growth and is particularly important for proliferation through ERK. This mechanism has been observed in other cancer types, further underscoring the diverse anticancer properties of VitC (Lee et al., 2017; Sinha et al., 2017; Wang et al., 2017; Yang et al., 2017).

Accumulating evidence shows the therapeutic potential of high-dose VitC in both preclinical and clinical settings (Böttger et al., 2021). Importantly, high-dose VitC has been widely shown to sensitize cancer cells to many conventional chemo- and radiation therapies and more recently to enhance cancer immunotherapy effects in multiple cancer types *in vivo* (Magrì et al., 2020). In the clinical setting, the use of high-dose VitC treatment, either alone or in conjunction with other therapies such as chemo- and/or radiotherapy, can produce notable anti-cancer effects, as shown in phase I/II studies and case reports. Especially in the palliative setting, the highly safe and tolerable high-dose VitC profile makes it a promising alternative for patients who typically have limited treatment options and a poor prognosis. Finally, it has been shown to significantly reduce the side effects caused by standard therapies in breast cancer and bone metastasis patients (Günes-Bayir and Kiziltan, 2015; Mansoor et al., 2021).

Large-scale pharmacological testing in conjunction with global profiling is of high interest to more fully understand the biology associated with sensitivity to high-dose VitC. System-wide approaches such as transcriptomic and proteomic studies can help capture the complexity of various cellular signalling pathways in an unbiased manner. Current transcriptome and proteome studies of high-dose VitC treatment are limited to a few small scale cell line studies (Aumailley et al., 2021; Böttger et al., 2021; Ngo et al., 2019). To our knowledge, this study is the first to perform large scale pharmacological analysis coupled to (phospho-)proteomics profiling. We report the sensitivities of VitC in 51 cell lines across 7 major cancer types that are all in the clinically achievable range, and identify proteins and pathways associated with VitC sensitivity. To further elucidate the mechanism of action of VitC, we profiled the (phospho-)proteomes of four PDAC cell lines treated with VitC at different doses and time points. Our data support further exploration of high dose VitC in the clinical setting.

## 2. MATERIALS AND METHODS

### 2.1. Cell lines and culture conditions

A total of 51 immortalized human cancer cell lines representative of pancreatic (PDAC), lung (LUAD, lung adenocarcinoma; LUSC, lung squamous cell carcinoma and SCLC, small cell lung cancer), prostate (PC), breast (BC) and colorectal (CRC) cancer were used in this study. Cells were cultured with Dulbecco’s modified Eagle’s medium (DMEM) or Roswell Park Memorial Institute (RPMI) 1640 Medium following manufacturers recommendations, and supplemented with 10% fetal calf serum (FCS) (Biowest) and 1% penicillin/streptomycin (P/S). For PC, LUSC and SCLC, all cell lines of the respective panel were cultured in RPMI medium. For BC, all cell lines of the respective panel were cultured in DMEM. For our 3 largest panels, LUAD, PDAC and CRC, different cell lines were cultured in different base media. For LUAD, this was RPMI for cell lines H2228, H1975, H522, H1650 and H322, and DMEM for A549 and SW1573. For PDAC, this was RPMI for cell lines CFPAC1, AsPC1, HPAC, BxPC3, PANC1 and Suit-2, and DMEM for cell lines Hs 766T, Panc-89, Capan-1, Capan-2, Mia PaCa-2 and PL45. For CRC, this was RPMI for cell lines LS-1034, LS-513, KM12, HCT116, COLO320-HSR and COLO205, and DMEM for cell lines RKO, SW948, SW48, HT55, HT29, SW1398 and SW620. All cell lines were cultured at 37°C in 5% CO2.

### 2.2. Preparation of 1M VitC stock solution

The 1M stock solution of VitC was freshly made for each experiment. VitC (05878 Fluka, Honeywell) was dissolved in type 1 MilliQ water (18 MW). The pH was then adjusted with 10M and 1M sodium hydroxide solution to pH 7.0-7.4. After adjusting the pH, the solution was filtered using a sterile syringe filter coupled to a filter unit with a low protein binding Durapore PVDF membrane (Merck Millipore). Finally, the solution was poured into an amber tube to protect from light exposure and sealed with parafilm to prevent interaction with oxygen.

### 2.3. Determination of cell line VitC sensitivity

Cells were trypsinized with 10x diluted Trypsin-EDTA (0.5%) (Gibco) solution for 5 minutes until detachment. Once trypsin was removed by centrifugation, cells were subsequently counted by using the LUNA-II™ Automated Cell Counter. Cells were then seeded in 96 well flat bottom Eppendorf plates at pre-determined densities. After 24 hours, allowing a complete adherence of the seeded cells, fresh 1M stock of VitC was prepared.

Forty out of fifty-four cell lines were then dosed per cell by counting them prior to treatment. This was done to adjust concentrations to pmol/cell based on their doubling time to ensure reproducibility as shown in previous studies (Schoenfeld et al., 2017). Average number of cells in the well prior to treatment (X) and expected number of cells after 24h (Y) were then used to adjust pmol/cell concentrations using the following formula: VitC concentration pmol/cell = X cells in 100 µL / Y * treatment volume. Cells were then treated for 2h with pharmacologic concentrations of VitC in a range of 0-20 mM. Respective dilutions were made with the corresponding media. After treatment, medium was carefully removed and replaced with fresh medium. A total of eleven cell lines were not dosed per cell but were included in the analysis using standard mM dosing.

Cell growth was determined by Sulforhodamine B (SRB) assay (Keepers et al., 1991; Vichai and Kirtikara, 2006) after 72 hours of treatment at 37°C and 5% CO2. In brief, a total of 25 μl of cold 50% Tri-chloroacetic acid (TCA) were added to each well and incubated for at least 60 min at 4°C. Plates were then washed five times with demi water and dried at room temperature (RT). 50 μl of SRB solution (0.4 % Sulforhodamine B (Sigma-Aldrich) and 1% acetic acid (Thermo Fisher Scientific) were subsequently added per well and cells were stained for 15 min at RT. Plates were then washed four times with 1% acetic acid solution and dried at RT. Finally, 150 μl of 10 mM Tris solution were added per well and plate was vortexed for 5 min. Data were read out by a Microplate Reader (Biotek, Synergy HT) at 490/540 nm. VitC sensitivity of all cell lines was determined by calculating the IC50s using pmol/cell and mM units. Since the SCLC cell line panel included suspension and semi-adherent cell lines, cell viability was determined using CellTiter-Fluor instead of SRB assay. Dose-response curves were generated and IC50 values calculated using GraphPad Prism software.

For catalase assay, cells were co-exposed with catalase (50U/ml) and VitC for 2h. Catalase was added 1 min prior to exposure as previously shown (Schoenfeld et al., 2017). Confluence images were taken by the Essen Bioscience IncuCyte™ ZOOM system and software.

### 2.4. Determination of cell line cisplatin sensitivity

Viability assays were performed to determine the IC50 values of cisplatin for the LUAD cell line panel. Depending on each cell lines’ proliferation time, 3000-5000 cells were seeded in 96 well flat bottom Eppendorf plates, left to attach for 24h and subsequently treated with a range of different cisplatin concentrations (0-1000 μM; Accord Healthcare) for 72h. Cell growth was determined using the SRB assay. Dose-response curves were generated and IC50 values calculated using GraphPad Prism software.

### 2.5. Intracellular ROS levels

Intracellular ROS levels were measured in Capan-2 and PANC-1 cell lines after 10 minutes, 2h and 24h of VitC treatment by using the OxiSelect™ Intracellular ROS Assay Kit (Green Fluorescence). Fluorescence was measured with a Biotek plate reader at 480 nm/530 nm wavelengths.

### 2.6. Preparation of untreated (baseline) cell lysates

Cells were grown up to ∼70% confluency in 10 cm petri dishes. Plates were washed with PBS and subsequently lysed in 1x SDS PAGE sample buffer. If the lysate was too viscous after scraping, 50-100 μL additional 1x SDS PAGE sample buffer was added. Next, cell lysates were sonicated using a Branson Digital Sonifier for 1 min. in 3 cycles of 20” on/20” off and then centrifugated for 10 min at max. speed. The supernatant was transferred to a new Eppendorf tube and stored at -20°C.

### 2.7. Preparation of on-treatment PDAC cell line lysates

For on-treatment (phospho-)proteome profiling, four PDAC cell lines (Suit-2, PANC-1, Capan-2 and Hs 766t) were seeded in 15 cm dishes and cultured until 60% confluency. Then, cell lines were treated with respective VitC IC50 doses (0.1, 0.14, 0.6 and 2 mM, respectively), and lysates were collected in a time series of 2, 4 and 24 hours post-treatment. For each time point and cell line, a control with no treatment was included (Fig. 9E). Furthermore, each condition consisted of two technical replicates, except for 2h- and 4h-control dishes, where each time point had one replicate due to the short time difference. After 2h of treatment, the media of the 4h and 24-hour dishes was refreshed with media without VitC, as previously described (Buranasudja et al., 2019; Michels and Frei, 2013). This was done to maintain the influx of H2O2 and to reduce the influence of breakdown products of VitC.

For each time point and cell line, we isolated the phosphoprotein fraction as previously described (Vallés-Martí et al., 2023). In brief, the supernatant was discarded and cells were washed with cold PBS at least three times. Immediately after, urea lysis buffer (9 M urea, 20 mM HEPES pH 8.0, 1 mM sodium orthovanadate, 2.5 mM sodium pyrophosphate, 1 mM β-glycerophosphate) was added and cells were removed from the substrate with a cell scraper. Lysate was collected in a 15 ml tube and stored at -80°C until sonication. The lysates were then subjected to sonication (3 cycles of 20” on/20” off at 18-micron amplitude) using a MSE Soniprep 150 sonicator and centrifugation (24,000G, 15 minutes, 10 ⁰C). Supernatant was transferred to a new tube, aliquoted and stored at 80°C until phosphopeptide capture. To quantify protein concentration, the Pierce BCA Protein Assay kit (Thermo Scientific) was used.

### 2.8. 5-band in-gel digestion

To determine optimal and comparable loading amounts for each untreated (baseline) cell line, different amounts of protein lysate were first loaded onto 12.5% SDS-PAGE test gels and proteins separated using the BioRad SDS-PAGE system (Biorad, Hercules, CA). Gels were run at 150 V for 50 min., fixed in 50% ethanol/3% phosphoric acid solution and stained with Coomassie brilliant blue G-250. Relative lysate amounts were quantified using ImageJ software. Subsequently, protein lysates were loaded onto a precast 4-12% gradient SDS-PAGE gel from the NuPAGE SDS-PAGE system (Invitrogen, Carlsbad, CA). The gel was run at 200 V for 40 min., fixed in 50% ethanol/3% phosphoric acid solution and stained with Coomassie brilliant blue G-250. Subsequent in-gel digestion was performed as described before (Piersma et al., 2010). In brief, proteins were reduced using 10 mM DTT and alkylated using 50 mM iodoacetamide. Then, in a laminar flow cabinet, each gel lane was cut into 5 bands, and each of them cut into ∼ 1 mm3 cubes, which were transferred to a pre- labelled Eppendorf tube. Every band was then separately digested overnight using trypsin. The peptides extracted from each gel band were subsequently stored at -20°C until LC-MS/MS analysis.

### 2.9. Sample preparation for phosphoproteomics

All lysates were reduced, alkylated and digested with Sequencing Grade Modified Trypsin (Promega, USA). For digestion and desalting an equivalent of 500 µg protein was used. Upon lysis proteins were reduced with DTT (dithiothreitol, 5 mM) and alkylated with IAA (iodoacetamide, 10 mM) followed by dilution to 2 M urea. Samples were digested with trypsin overnight (1:100 m/m) and then acidified by addition of 0.1% Tri-fluoroacetic acid (TFA). The digests were desalted using Oasis HLB columns (500 mg capacity, Waters), eluted in 0.1% TFA, 80% acetonitrile (ACN) and lyophilized for 48 h in a freeze dryer (Christ alfa 2-4 LSC).

Phosphopeptide enrichment was performed by IMAC (immobilized metal ion affinity chromatography), using the BRAVO AssayMap liquid handling robot (Agilent) as previously described (Vallés-Martí et al., 2023). In brief, 200 µg peptides for each sample were acidified to 0.1% TFA, and applied to a 10 mg OASIS HLB cartridge (Waters) previously activated with acetonitrile and equillibrated with 0.1% TFA. After washing the columns twice with 0.1% TFA, the bound phosphopeptides were eluted in 100 µl 5% NH4OH/30% ACN and then transferred to glass-lined autosampler vials and dried in a vacuum centrifuge at 45 °C. Finally, samples were re-dissolved in 20 µl 0.5% TFA/4% ACN prior to injection and stored at 4 °C until LC-MS/MS analysis. Two replicates of the control lysate of HCT116 (colon carcinoma cell line) were included in the processing as workflow control.

### 2.10. LC/MS-MS Analysis

For the untreated (baseline) proteomic profiling, peptides were separated by an Ultimate 3000 nanoLC-MS/MS system (Dionex LC-Packings, the Netherlands) equipped with a 40 cm × 75 μm ID fused silica column custom packed with 1.9 μm 120 Å ReproSil Pur C18 aqua (Dr Maisch GMBH, Germany). After sample injection, peptides were trapped at 6 μl/min on a 10 mm × 100 μm ID trap column packed with 5 μm 120 Å ReproSil Pur C18 aqua at 2% buffer B (buffer A: 0.5% acetic acid (Fischer Scientific), buffer B: 80% ACN, 0.5% acetic acid) and separated at 300 nl/min in a 10–40% buffer B gradient in 90 min (120 min inject-to-inject) at 35 °C. The eluting peptides were ionized at a potential of + 2 kVa into a Q Exactive mass spectrometer (Thermo Fisher, Germany). Intact masses were measured at resolution 70.000 (at m/z 200) in the Orbitrap using an AGC target value of 3 × 106 charges. The top 10 peptide signals (charge-states 2+ and higher) were submitted to MS/MS in the HCD (higher-energy collision) cell (1.6 amu isolation width, 25% normalized collision energy). MS/MS spectra were acquired at resolution 17.500 (at m/z 200) in the orbitrap using an AGC target value of 2 × 105 charges and an underfill ratio of 0.1%. Dynamic exclusion was applied with a repeat count of 1 and an exclusion time of 30 s.

For the on-treatment phosphoproteomic profiling, peptides were separated on an Ultimate 3000 nanoLC-MS/MS system (Dionex LC-Packings, Amsterdam, the Netherlands) equipped with a 50-cm 75 µm ID C18 Acclaim pepmap column (Thermo Scientific). After injection, peptides were trapped at 3 μl/min on a 10-mm, 75-μm ID Acclaim Pepmap trap column (Thermo Scientific) in buffer A (buffer A: 0.1% formic acid, buffer B: 80% ACN/0.1% formic acid), and separated at 300 ml/min with a 10–40% buffer B gradient in 90 min (120 min inject-to-inject). Eluting peptides were ionized at a potential of +2 kV and introduced into a Q Exactive HF mass spectrometer (Thermo Fisher, Bremen, Germany). Intact masses were measured in the orbitrap with a resolution of 120,000 (at m/z 200) using an automatic gain control (AGC) target value of 3 × 106 charges. Peptides with the top 15 highest signals (charge states 2+ and higher) were submitted to MS/MS in the higher-energy collision cell (1.6-Da isolation width, 25% normalized collision energy). MS/MS spectra were acquired in the Orbitrap with a resolution of 15k (at m/z 200) using an AGC target value of 1 × 106 charges and an underfill ratio of 0.1%. Dynamic exclusion was applied with a repeat count of 1 and an exclusion time of 30 s.

### 2.11. Phosphoprotein identification

MS/MS spectra were searched against the Swissprot human reference proteome 2020_04_cannonical_and_isoform.FASTA (42347 entries) using MaxQuant (Cox and Mann, 2008) 1.6.10.43. Enzyme specificity was set to trypsin and up to two missed cleavages were allowed. Cysteine carboxamidomethylation (+ 57.021464 Da) was treated as fixed modification and methionine oxidation (+ 15.994915 Da) and N-terminal acetylation (+ 42.010565 Da) as variable modifications. Peptide precursor ions were searched with a maximum mass deviation of 4.5 ppm and fragment ions with a maximum mass deviation of 20 ppm. Peptide and protein identifications were filtered at an FDR of 1% using the decoy database strategy. The minimal peptide length was 7 amino-acids and the minimum Andromeda score for modified peptides was 40 and the corresponding minimum delta score was 6. Proteins that could not be differentiated based on MS/MS spectra alone were grouped to protein groups (default MaxQuant settings). Protein expression searches were performed with the label-free quantification option selected, and match between runs option was off. Proteins were quantified using spectral count MS data, which was subsequently normalized by a global normalization to the total count of each sample.

MS/MS spectra of IMAC phosphopeptide enrichment experiments were searched separately against the Swissprot human FASTA file (canonical and isoforms; 2021, 42383 entries) using MaxQuant 1.6.10.43. Enzyme specificity was set to trypsin, and up to two missed cleavages were allowed. Cysteine carboxamidomethylation (+57.021464 Da) was treated as fixed modification and serine, threonine, and tyrosine phosphorylation (+79.966330 Da), methionine oxidation (+15.994915 Da), and N-terminal acetylation (+42.010565 Da) as variable modifications. Peptide precursor ions were searched with a maximum mass deviation of 4.5 ppm and fragment ions with a maximum mass deviation of 20 ppm. Peptide and protein identifications were filtered at a false discovery rate of 1% using a decoy database strategy. The minimal peptide length was set at 7 amino acids, the minimum Andromeda score for modified peptides was 40, and the corresponding minimum delta score was 6. Proteins that could not be differentiated based on MS/MS spectra alone were clustered into protein groups (all default MaxQuant settings). Phosphopeptide identifications were propagated across samples using the ‘match between runs’ option checked. For phosphopeptide data, we used data from the MaxQuant ‘modificationSpecificPeptides’ table. For phosphosite data, we used data from the MaxQuant ‘Phospho (STY) Sites’ table. Phosphopeptides and phosphosites were quantified from the area under the curve of the MS1 signal of each eluting peptide (‘Intensity’ in MaxQuant). Human Swiss-Prot database used for raw data search was acquired from the UniProt database (https://www.uniprot.org/). For calculation of kinase INKA scores, phosphopeptide MS/MS spectral counts were calculated from the MaxQuant evidence file using R.

### 2.12. Statistical analysis and pathway annotation

To unravel biology and proteins associated to VitC sensitivity, and considering the unique molecular characteristics of each cancer type, we performed two complementary statistical analyses per panel: a Spearman correlation and a two-group comparison. Spearman correlation analyses were performed between VitC half-maximal inhibitory concentrations (IC50s) and the protein expression for the whole panel, using the R package *stats*. Group comparisons with proteomic data (2 lowest IC50 vs 2 highest IC50, or 4 vs 4 cell lines, depending on panel size) were performed by bb test using the *ion* package in R. For proteins correlating with VitC IC50 values, a filtering threshold of rho ≥ 0.7 and a *pvalue* of ≤ 0.05 was the standard for the majority of panels, except for CRC for which the rho threshold was set to 0.6 to increase number of identifications. Two group comparison analyses were performed on the cell lines with lowest and highest VitC sensitivity per panel, whereby number of samples in low and high sensitivity groups differed per panel depending on panel size. Filtering threshold was set to a fold change of ≥ 2 and a *pvalue* of ≤ 0.05. The overlap between the two statistical analyses was performed on the significant candidates associated with low and high sensitivity, and the enriched biology was further investigated using gProfiler (Raudvere et al., 2019). Top 10 enriched and non-redundant biology terms were selected. A more stringent analysis was followed to select top 20 proteins associated with high and low sensitivity, including only significant hits which had a rho threshold of ≥ 0.7 and a fold change of ≥ 1.5. Top candidates involved in known VitC associated processes (Böttger et al., 2021; Ngo et al., 2019) were further annotated, as well as those that were only significant in one analysis but involved in these processes. To illustrate common enriched biology associated with VitC sensitivity among cancer cell line panels, the top50 significant (*adjusted pvalue* < 0.05) gProfiler output terms (GO:BP and GO:CC) were overlapped between all 7 panels using Venny v2.1 (Oliveros, J.C., 2015) . All shared terms were then summarized by Revigo v1.8.1 using SimRel (default) semantic similarity measure and small (0.5) resulting list (Supek et al., 2011). Separate analyses were performed for terms associated with high and low sensitivity.

For gene set enrichment analysis (GSEA), significant proteins were ranked by multiplying the sign of the fold change by the -log10 *pvalue*. This was used as input for GSEA (Subramanian et al., 2005) using the *fgsea* (Sergushichev, 2016) package in R. Gene signatures were retrieved with the *msigdbr* (Liberzon et al., 2011) package. Single sample gene set enrichment analysis (ssGSEA) was performed using the *GSVA* R package (Hänzelmann et al., 2013). For the targeted analysis, known processes associated with VitC sensitivity (Böttger et al., 2021; Ngo et al., 2019) were retrieved from Reactome (Gillespie et al., 2022) and Gene Ontology (Ashburner et al., 2000; Consortium et al., 2023) of biological and molecular function processes.

For the phosphoproteome data analyses, we used the log-transformed median-site normalized intensities. To determine differences between VitC and control samples, fold changes (FC) were subsequently calculated with the normalized intensities. In brief, averaged VitC treated replicates were divided by the averaged controls per timepoint to calculate upregulation, and -1 times the averaged controls divided by averaged VitC treated replicates per timepoint was used to calculate downregulation. These calculations resulted in FC where 1 and -1 were equivalent to no change between VitC treated samples and controls. Kinase activity was inferred from phosphoproteomic data with the Integrative iNferred Kinase Activity (INKA) tool as previously described by Beekhof et al. (Beekhof et al., 2019) and available at https://inkascore.org/. The kinases were ranked according to the sum of all INKA scores per kinase of all samples divided by the number of samples. Based on the ranking, the top 20 ranked kinases and boxplots of the INKA scores were made. Fold changes were then calculated per cell line and timepoint with the same method as described above and heatmaps with the corresponding FC were made per cell line.

## 3. RESULTS

### 3.1. High-dose VitC pan-cancer sensitivities

To further underscore the potential of high-dose VitC as a broadly acting anti-cancer agent, we determined VitC responses in a large cancer cell line panel containing seven major cancer types (PDAC, pancreatic ductal adenocarcinoma; CRC, colorectal cancer; LUAD, lung adenocarcinoma; LUSC, lung squamous cell carcinoma; SCLC, small cell lung cancer; PC, prostate cancer; BC, breast cancer; Fig.1A).

**Fig. 1.**
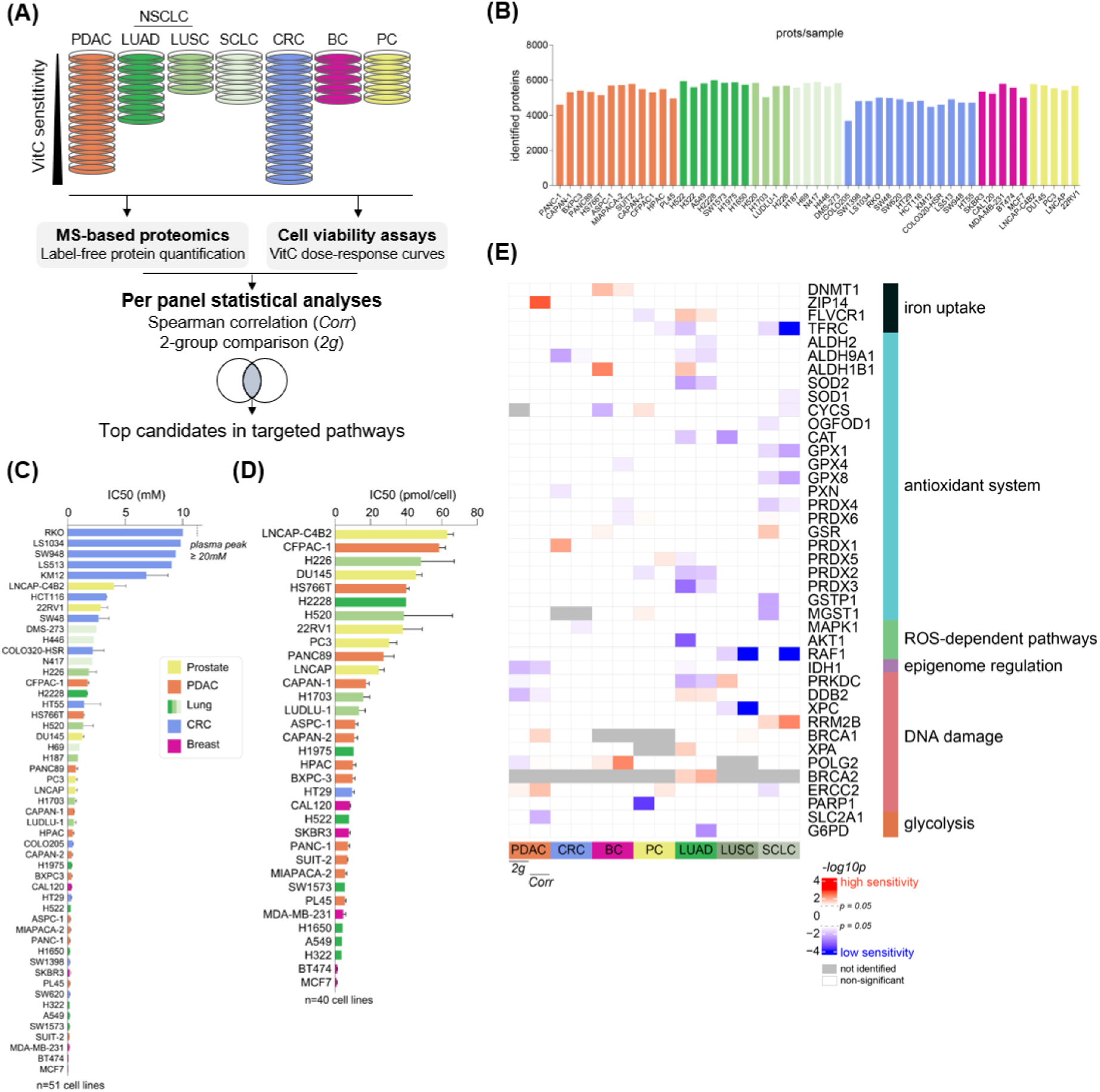
Pan-cancer proteomic analysis of VitC sensitivity. (A) Study design of the pan-cancer proteome profiling of 51 human cell lines divided by cancer (sub)type panel: PDAC, LUAD, LUSC, SCLC, CRC, BC and PC, with the determination of VitC sensitivities. The candidate selection of sensitivity-associated proteins per panel was supported by two complementary statistical analyses: a Spearman correlation of protein expression with IC50 values (Corr), and a two-group comparison between protein expression in more and less sensitive cell lines (2g). (B) Number of identified proteins per sample. (C) (D) VitC sensitivities plotted as IC50s, measured by cell viability assays and represented (C) in millimolar (n=51 cell lines) and (D) in pmol/cell range (n=40 cell lines). Mean ± SEM plotted, average n ≥ 2. (E) Targeted analysis of potential VitC-associated processes (Böttger et al., 2021). The significance by two group comparison and Spearman correlation is shown per panel. Proteins with at least one significant *pvalue* in at least one panel were included.

Pharmacological VitC testing was performed in all cell lines, which were treated with ten VitC concentrations (0-20 mM), and the respective sensitivity was inferred by using IC50s at mM and pmol/cell dosing (Fig. 1C-D, respectively). For the majority of cell lines, a 5 mM VitC dose was sufficient to inhibit 50% of cell growth, and among all, average IC50 was 1.7±0.4 mM (range 0.036-10.02 mM). Due to the known VitC and H_2_O_2_ toxicity dependency on cell density (Doskey et al., 2015; Schoenfeld et al., 2017), a dosing per cell metric scheme was applied for a selection of cell lines (n=40 in PDAC, LUAD, LUSC, PC and BC panels) and an average of 18.7±3 pmol/cell (range 0.95-63.2 pmol/cell) was observed (Fig. 1D). Interestingly, the two metrics -mM and pmol/cell-showed comparable results to a large extent (Fig. 1C-D, Pearson coefficient R=0.892, *pvalue*=1.143e-13).

Altogether, our large scale pan-cancer pharmacological analysis showed that all cancer cell lines were killed at clinically achievable concentrations (below two-fold the 20 mM plasma peak), extending the finding of previous smaller scale cancer cell line studies (Böttger et al., 2021; Chen et al., 2005) and underscoring its promise as a broad anti-cancer therapy.

### 3.2. Per cancer cell line panel analysis of biological processes and proteins associated with high-dose VitC sensitivity

Little is known about the proteome association with high-dose VitC cancer vulnerabilities. In fact, to our knowledge, the present study is the first to evaluate the baseline proteomic features associated with high-dose VitC responses in a large pan-cancer panel. To this end, polyacrylamide gel fractionation coupled to MS-based label-free proteomics was applied to the 51 cancer cell lines for which we determined the VitC IC50 values. On average, 5352 proteins were identified per sample (Fig.1B). Because these cell line panel datasets were generated at different points in time, we analysed the data per panel.

First, we evaluated the expression of several proteins of processes previously found to be associated with VitC sensitivity such as iron uptake, antioxidant defence system, DNA repair or glucose metabolism (Böttger et al., 2021; Ngo et al., 2019) (Fig. 1E). Two complementary statistical approaches were applied to each panel and proteins significant in either statistical analyses, 2-group comparison or Spearman correlation, were evaluated. As expected, proteins involved in the antioxidant system, such as ROS scavengers CAT, SOD1/2, GPXs and PRDXs, were significantly increased in less sensitive cell lines from multiple cancer types, particularly LUAD and SCLC. Similarly, ROS-related pathways (Kirtonia et al., 2020) were found high in less sensitive cell lines. Increased intracellular iron, in the Fe^2+^ form, reacts with hydrogen peroxide to form hydroxyl radicals, thus damaging DNA, proteins and lipids, and leading to cancer cell death, thereby increasing VitC sensitivity. Iron transporter TFRC expression, that mediates transport of iron bound to transferrin, was found high in less sensitive PC, LUAD and SCLC cell lines. However, proteins involved in “free” iron transport and mobilization (Brown et al., 2020; Lane and Richardson, 2014), e.g. DNMT1, ZIP14 and FLVCR1, were found to be significantly increased in sensitive cell lines, especially in the BC, PDAC and LUAD panels. Increased abundance of glucose transporter SLC2A1 (GLUT1), which can transport the oxidized form of VitC into the cell (Vera et al., 1993), was found to be significant in less sensitive PDAC cell lines. Glycolysis-related protein G6PD, which is also an oxidative stress protector, was also found significantly high in less sensitive LUAD cell lines. On the contrary, DNA damage/repair proteins such as DDB2, RRM2B, BRCA1, XPA and POLG2 were found to be particularly elevated in cells with high sensitivity, which may be also linked to proliferation signals. IDH1, regulator of DNA methylation, was high in less sensitive cell lines in the PDAC and LUAD panels. For functional data mining and selection of VitC sensitivity-associated proteins, the intersection of the 2-group and correlation statistical analyses was used. Below we present the results per cancer panel.

For one of the deadliest cancer types, PDAC, we investigated a large panel containing 12 PDAC cell lines, which displayed a diverse sensitivity range (5.5-58.6 pmol/cell and 0.12-1.72 mM, Fig. 2A-C). Of note, no association between sensitivity and the epithelial/mesenchymal phenotypes was found (Fig. 2A), which underscores the one-size-fits-all capability of VitC as an anti-cancer agent. Proliferative functions such as cell cycle, RNA metabolism, translation and mitochondrial processes were enriched in highly sensitive cell lines (Fig. 2D). High-sensitivity top 20 candidates were found to be mainly involved in RNA splicing (e.g. SNRNP200, SRRM2 and AQR) and RNA translation and transcription (e.g. GTF3C1, RPS5, RPS7, RRP12, SIN3A, BMS1, GTF3C5 and SMG1). In addition, nuclear pore complex protein TPR, which is also involved in glucokinase transport, and protein repair protein PCMT1 were among the top 20 high sensitivity candidates (Fig. 2F). Cells with lower sensitivity displayed higher cell adhesion, extracellular vesicle, and mevalonate pathway functions, the latter being involved in cholesterol synthesis and protein prenylation (Fig. 2E). Among the top 20 candidates associated with lower VitC sensitivity were the superoxide producer CYBA, DNA damage repair proteins UBE2V1, PPP5C and POLB, calcium-ion binding proteins S100A6, S100A16, S100A13 and CIB1, the cholesterol synthesis protein MVD, and two important glycolysis and TCA proteins, PFKL and MUT, respectively (Fig. 2G).

**Fig. 2.**
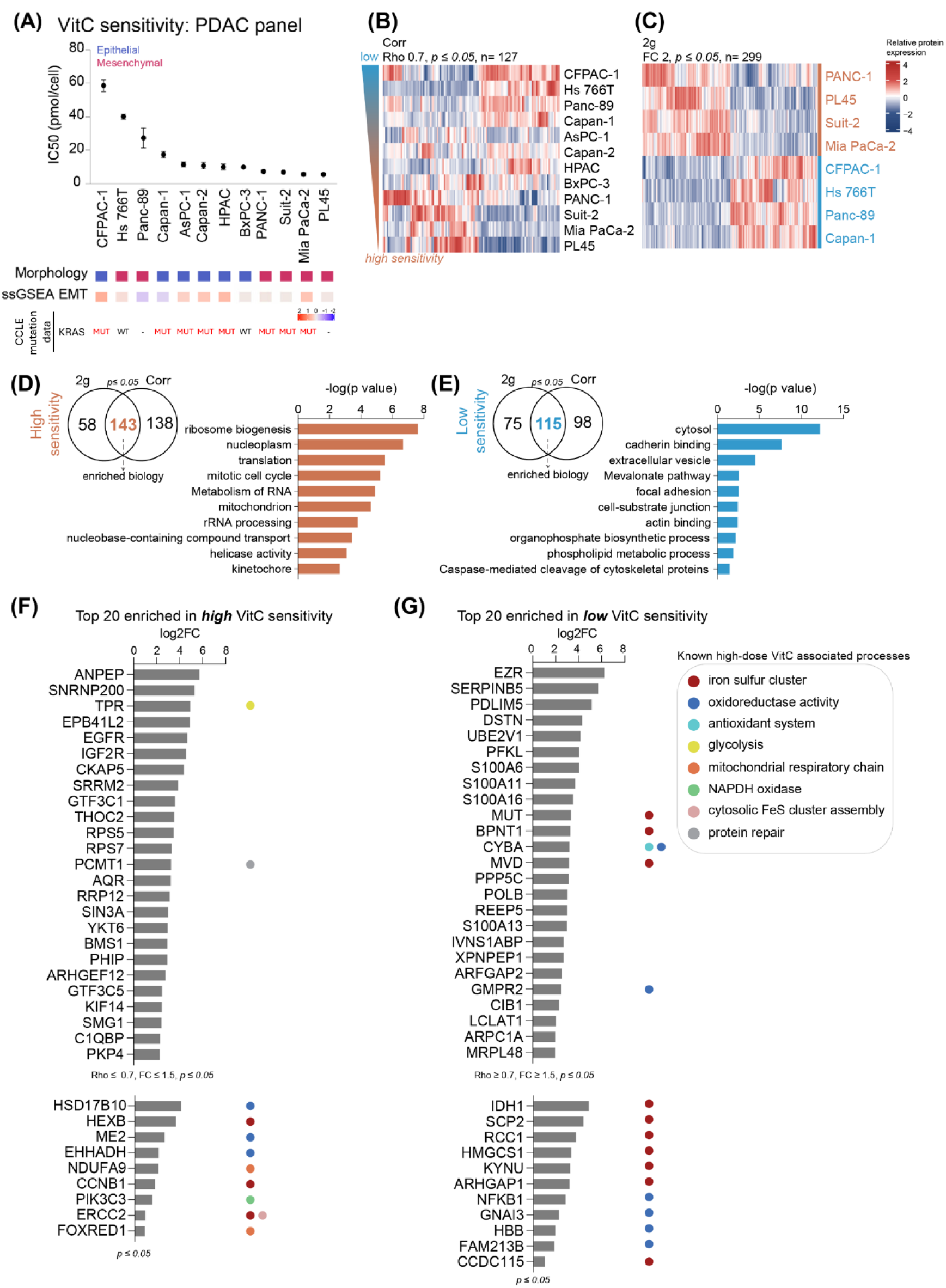
Pancreatic ductal adenocarcinoma VitC sensitivity and protein biomarker discovery analysis. (A) VitC sensitivities (IC50 pmol/cell) of n=12 PDAC cell lines. Epithelial and mesenchymal classification is annotated based on morphology and ssGSEA EMT signature. KRAS mutation status annotations were retrieved from CCLE mutations database. WT, wild type; MUT, somatic driver mutation at G12; -, not represented in CCLE. (B) Hierarchical clustering heatmap of proteins correlated with high and low VitC sensitivity (Spearman correlation rho ≥ or ≤ 0.7, *pvalue* ≤ 0.05, n=127 identified proteins). (C) Hierarchical clustering heatmap of proteins enriched in highly sensitive and less sensitive cell lines (2-group comparison, FC ≥ or ≤ 2, *pvalue* ≤ 0.05, n=299 identified proteins). (D), (E) Venn diagram describing the overlap between significant proteins found in both statistical analysis, 2-group comparison and Spearman correlation for (D) high and (E) low VitC sensitivity, as input for the top 10 enriched biology terms. Terms for (D) high and (E) low VitC sensitivity are ranked on –*log10 pvalue*. (F), (G) Top 20 candidate markers associated with (F) high and (G) low VitC sensitivity ranked on log2FC. Gene ontology terms of known VitC related processes were further annotated. Significant proteins found in these VitC-related processes were also added.

Likewise, we investigated a large panel containing 13 colorectal cancer (CRC) cell lines, which displayed a sigmoid sensitivity range (0.16-10 mM, Fig.3A-C). CMS subtypes did not correlate with VitC sensitivity, confirming the universal anti-cancer effect of VitC, regardless of molecular subtype, as already observed in the PDAC panel. Processes associated with high VitC sensitivity were again linked to proliferation including mRNA metabolism and splicing, cell cycle and DNA damage responses (Fig. 3D). Interestingly, a great number of pre-mRNA splicing proteins were found among the top 20 candidates, such as SF3A1, LUC7L2, PUF60, TRA2B, PRPF38A and SRSF10, as well as transcription regulators HNRNPH1, CNBP, ATRX, EIF4H, RAE1 and SMARCD1 (Fig. 3F). Proteins involved in energy and glucose metabolism such as CKB, VDAC2, AKR1B1 and RAE1 were also indicators of high VitC sensitivity. Processes associated with lower sensitivity were vesicle-mediated transport, glucose metabolism, cell adhesion and protein transport (Fig. 3E). Interestingly, among the top 20 proteins were mitochondrial activity related proteins such as HSPA9, HADHA and STOML2, together with other metabolic proteins such as the key player of glycolysis, GPI, lipid synthesis protein ACSL1, and SPR, promoter of BH4, which is a potent radical-trapping antioxidant (Soula et al., 2020) (Fig. 3G). Another significant protein that could influence VitC sensitivity was pyruvate dehydrogenase complex member PDHB, involved in the glycolysis to TCA cycle step.

**Fig. 3.**
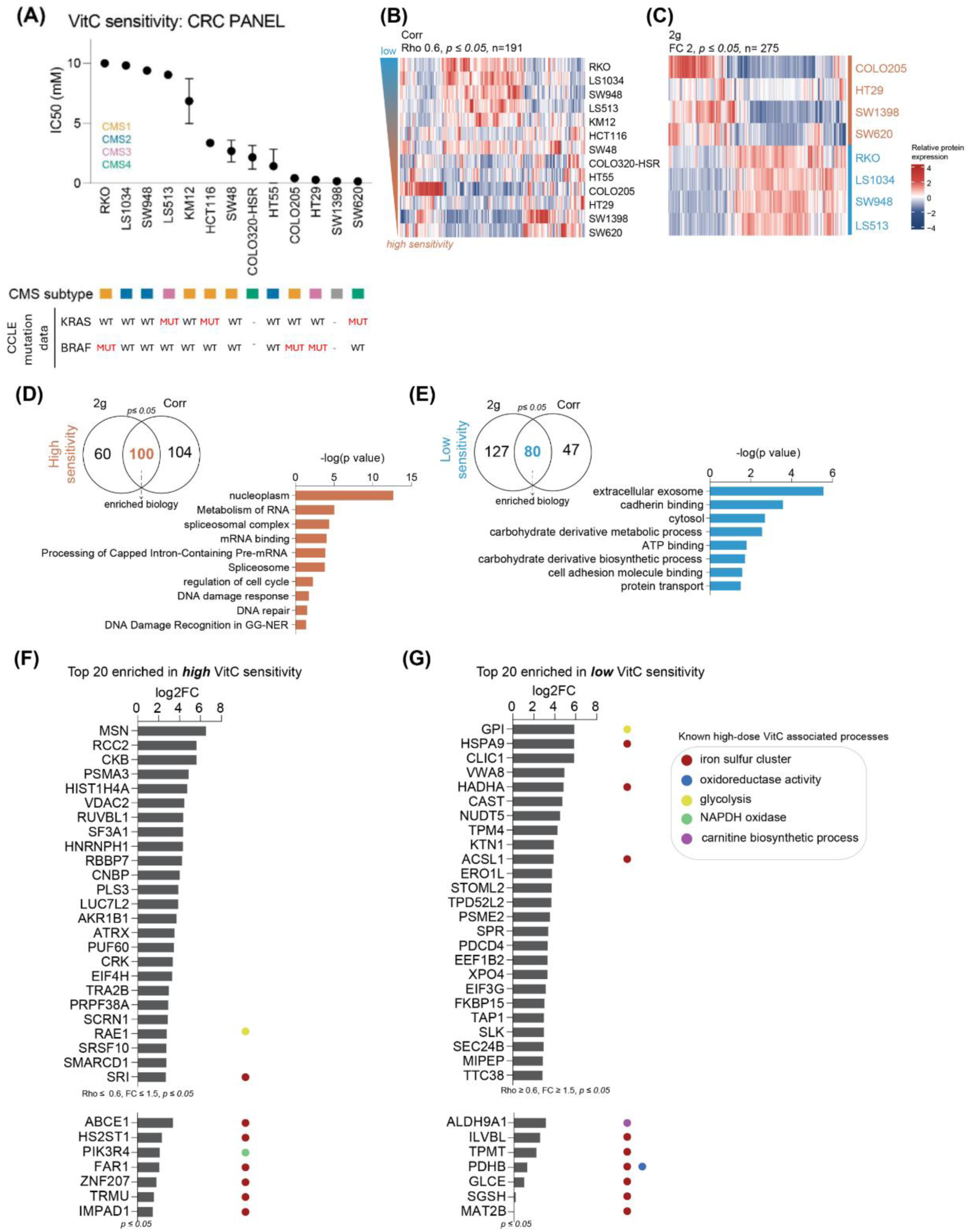
Colorectal cancer VitC sensitivity and protein biomarker discovery analysis. (A) VitC sensitivities (IC50 mM) of n=13 CRC cell lines. CMS subtype information was retrieved from two recent studies (Frejno et al., 2017; Linnekamp et al., 2018). KRAS and BRAF mutation status annotations were retrieved from CCLE mutations database. WT, wild type; MUT, somatic driver mutation at G12 or G13 (KRAS) and V600 (BRAF); -, not represented in CCLE. (B) Hierarchical clustering heatmap of proteins correlated with high and low VitC sensitivity (Spearman correlation rho ≥ or ≤ 0.6, *pvalue* ≤ 0.05, n=191 identified proteins). (C) Hierarchical clustering heatmap of proteins enriched in highly sensitive and less sensitive cell lines (2-group comparison, FC ≥ or ≤ 2, *pvalue* ≤ 0.05, n=275 identified proteins). (D), (E) Venn diagram describing the overlap between significant proteins found in both statistical analysis, 2-group comparison and Spearman correlation for (D) high and (E) low VitC sensitivity, as input for the top 10 enriched biology terms. Terms for (D) high and (E) low VitC sensitivity are ranked on –log10 *pvalue*. (F), (G) Top 20 candidate markers associated with (F) high and (G) low VitC sensitivity ranked on log2FC. Gene ontology terms of known VitC related processes were further annotated. Significant proteins found in these VitC-related processes were also added.

As for CRC, a sigmoid response was also observed for the breast cancer panel containing 5 cell lines, which showed the highest sensitivity among all panels (0.95-8.4 pmol/cell and 0.036-0.32 mM, Fig. 4A-C). All cell lines were thus considered to be relatively sensitive to VitC, regardless of breast cancer subtypes. Biological processes such as DNA replication and DNA damage responses, together with nuclear processes and telomere C-strand synthesis were enriched in the more sensitive cells (Fig. 4D). Top high-sensitivity candidate proteins were strongly involved in DNA repair, such as HIST1H2BM, DDB1, TOP2B, HIST2H3A, MDC1, SUPT16H and PDS5A, as well as in RNA splicing, as previously also observed in the PDAC and CRC panels, represented by proteins such as SFPQ, SF3B1, DHX15, DDX46 and THRAP3 in the breast cancer panel (Fig. 4F). Significant candidates involved in oxidative stress were also identified: OXR1, which regulates kinase signalling in response to stress, and QSOX2, a hydrogen peroxide generator, which supports the fact that sensitive cells indeed have boosted levels of ROS (Schoenfeld et al., 2017). Terms enriched in breast cancer cell lines with reduced VitC sensitivity were ribosomal and amino acid metabolism, mitochondrial and extracellular exosome (Fig. 4E). Ribosomal-related proteins, e.g. RPL5, RPL7, RPS6, RPL13A, MRTO4 and RPL13, were particularly represented in the top 20 candidates for low VitC sensitivity (Fig. 4G). In addition, we identified two-crucial proteins for the mitochondrial beta-oxidation pathway, ACAT1 and ECHS1. Interestingly, we also identified PYCR1, which promotes proline synthesis by promoting glycolysis and nucleotide synthesis, as well as inhibiting ROS levels (Tanner et al., 2018). Nucleotide metabolism-related proteins were also identified (ADK, AHCY and TK1). Two proteins were involved in oxidative stress homeostasis: PHB2, which has recently been shown to be involved in the regulation of mitochondrial ROS levels in glioblastoma (Huang et al., 2021), and GSTK1, which is an antioxidant contained in peroxisomes. Other significant and interesting proteins were also involved in oxidative stress homeostasis, e.g. LANCL1, which functions as a glutathione transferase, as well as ATP6V0D1, involved in iron homeostasis by triggering Fe^2+^ prolyl hydroxylases, which in turn hydroxylate HIF1A and target it to proteasomal degradation (Miles et al., 2017). We also identified four mitochondrial proteins, NDUFA9, NDUFS6, NDUFA5, NDUFAF5, which are part of mitochondrial respiratory chain Complex I, associated to low VitC sensitivity.

**Fig. 4.**
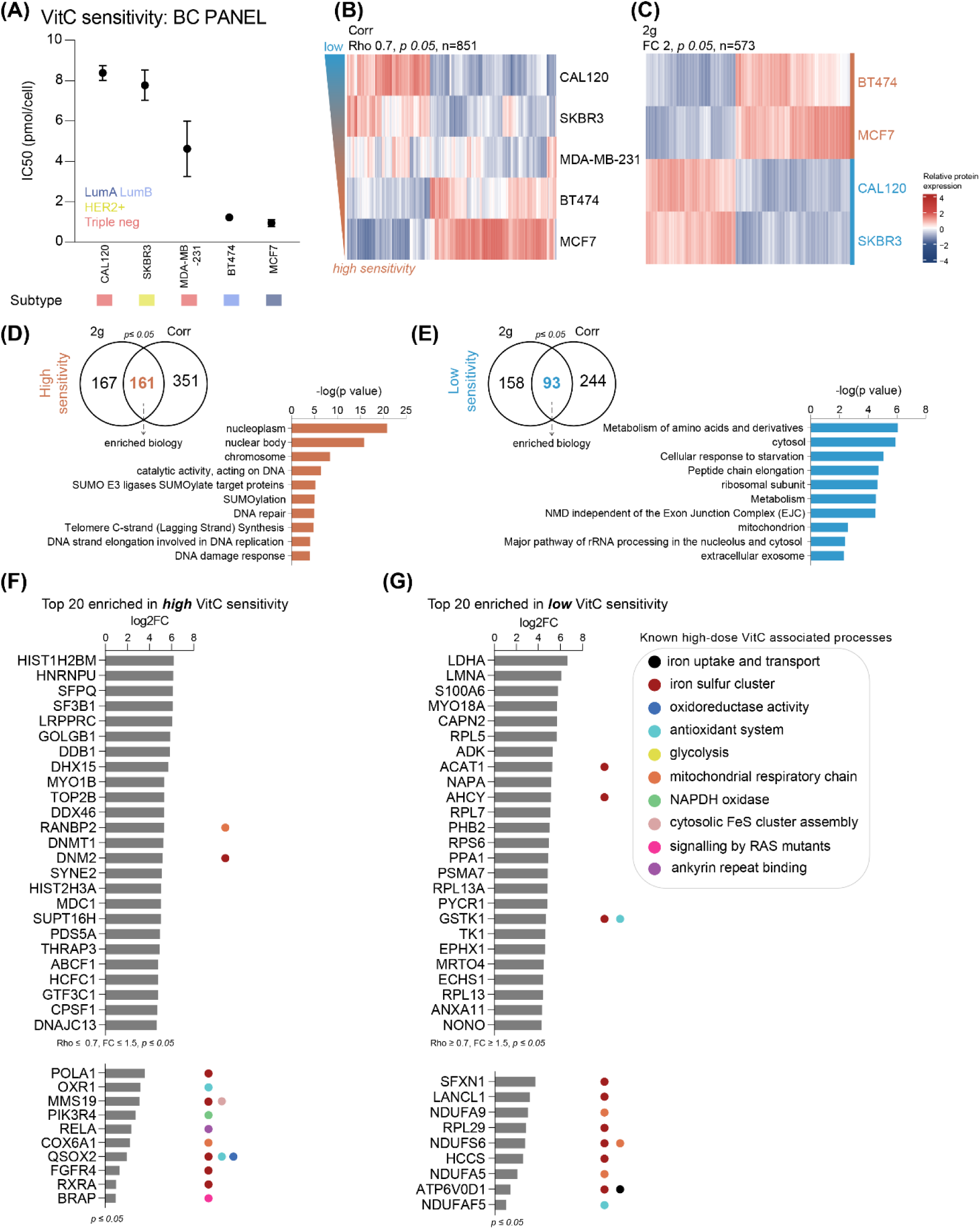
Breast cancer VitC sensitivity and protein biomarker discovery analysis. (A) VitC sensitivities (IC50 pmol/cell) of n=5 BC cell lines. Molecular subtype was annotated based on a reference study (Dai et al., 2017). (B) Hierarchical clustering heatmap of proteins correlated with high and low VitC sensitivity (Spearman correlation rho ≥ or ≤ 0.7, *pvalue* ≤ 0.05, n=851 identified proteins). (C) Hierarchical clustering heatmap of proteins enriched in highly sensitive and less sensitive cell lines (2-group comparison, FC ≥ or ≤ 2, *pvalue* ≤ 0.05, n=573 identified proteins). (D), (E) Venn diagram describing the overlap between significant proteins found in both statistical analysis, 2-group comparison and Spearman correlation for (D) high and (E) low VitC sensitivity, as input for the top 10 enriched biology terms. Terms for (D) high and (E) low VitC sensitivity are ranked on –log10 *pvalue*. (F), (G) Top 20 candidate markers associated with (F) high and (G) low VitC sensitivity ranked on log2FC. Gene ontology terms of known VitC related processes were further annotated. Significant proteins found in these VitC-related processes were also added.

Of note, prostate (Fig. S1), small cell lung (Fig. S2), lung adenocarcinoma (Fig. 5) and lung squamous cell (Fig. S3) cancer panels were analysed in the same manner. Interestingly, the prostate cancer panel behaved differently, with proliferation signals (e.g. cell cycle, telomere maintenance) being observed in the less sensitive cell lines, as opposed to in the more sensitive cell lines in all other cell line panels. Furthermore, lung cancer types and subtypes were overall quite sensitive but displayed different sensitivities: LUAD was the most sensitive panel (mean 0.42±0.56 mM), followed by LUSC (mean 1.1±0.62 mM) and SCLC (mean 1.8±0.75 mM). In line with this, we found increased SLC2A3 (GLUT3) in LUAD sensitive cell lines (Fig. 5F), which is involved in the transmembrane transport of dehydroascorbic acid, the oxidized form of VitC. Iron regulators FLVCR1 and HMOX2 were additionally increased in this panel (Fig. 5F-G), which confirms the crucial role of iron and VitC uptake in VitC sensitivity.

**Fig. 5.**
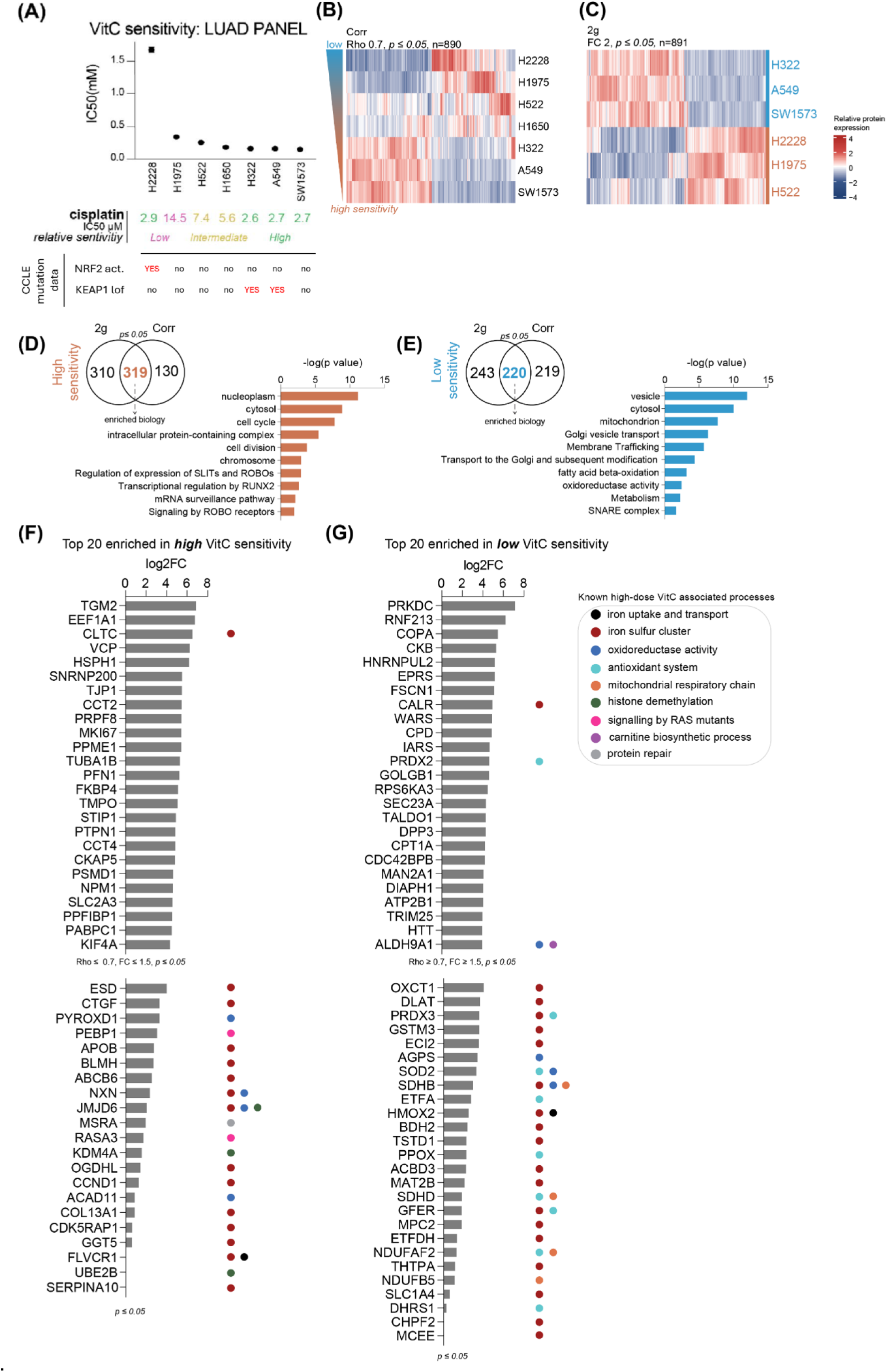
Lung adenocarcinoma VitC sensitivity and protein biomarker discovery analysis. (A) VitC sensitivities (IC50 mM) of n=7 LUAD cell lines. Cisplatin sensitivities (IC50s μM) were further annotated. NRF2 activating (act.) and KEAP1 loss-of-function (lof) mutation status annotations were retrieved from CCLE mutations database. (B) Hierarchical clustering heatmap of proteins correlated with high and low VitC sensitivity (Spearman correlation rho ≥ or ≤ 0.7, *pvalue* ≤ 0.05, n=890 identified proteins). (C) Hierarchical clustering heatmap of proteins enriched in highly sensitive and less sensitive cell lines (2-group comparison, FC ≥ or ≤ 2, *pvalue* ≤ 0.05, n=891 identified proteins). (D), (E) Venn diagram describing the overlap between significant proteins found in both statistical analysis, 2-group comparison and Spearman correlation for (D) high and (E) low VitC sensitivity, as input for the top 10 enriched biology terms. Terms for (D) high and (E) low VitC sensitivity are ranked on –log10 *pvalue*. (F), (G) Top 20 candidate markers associated with (F) high and (G) low VitC sensitivity ranked on log2FC. Gene ontology terms of known VitC related processes were further annotated. Significant proteins found in these VitC-related processes were also added.

### 3.3. Implications for combination treatment of chemotherapy-resistant lung cancer

Interestingly, among the top 5 proteins most enriched in highly VitC sensitive LUAD cancer cell lines were several candidates that have previously been implicated in chemotherapy resistance (TGM2 (Huang et al., 2015) and VCP (Costantini et al., 2021), Fig. 5F). Platinum-based chemotherapy is the central component of most treatment regimens administered to patients with NSCLC; however clinical benefit is often hampered by chemoresistance (Cortés et al., 2015; Michael et al., 2020). Previously, we have profiled the lysates and secretomes of a panel of NSCLC cell lines in relation to cisplatin IC50 values, and revealed biofluid biomarker candidates with potential predictive value in determining cisplatin response (Böttger et al., 2019). In the present study, we have extended the determination of cisplatin sensitivities to the entire LUAD panel. Strikingly, we found the least cisplatin sensitive cell line of the LUAD panel (H1975), as determined by assessing cell viability in response to different cisplatin doses, to be part of the more VitC sensitive cell line group. This further supports a role for VitC as a non-toxic alternative or combination treatment option for chemotherapy resistant lung cancer populations (Fig. 5A).

Incidentally, a similar observation was made for CRC (data not shown, as only a subset of our CRC panel was represented in the NCI-60 database), underscoring the point that cytotoxic agent- and VitC-dependent cell killing do not rely on the same mechanisms. In line with this conclusion, high-dose VitC has been reported to enhance the effectivity of a wide variety of chemotherapeutic drugs in various preclinical cancer cell type models, often in a synergistic manner or by enhancing treatment efficacy (Table 1 and Fig. 2 in (Böttger et al., 2021).

### 3.4. Evaluation of genomic associations to high-dose VitC sensitivity

Based on previous reports suggesting a possible link between KRAS/BRAF mutation and enhanced VitC susceptibility in CRC (Yun et al., 2015), we set out to explore how known oncogenic driver mutations correlate with VitC sensitivity. We annotated common somatic mutations that may influence VitC susceptibility in cell line panels of cancers in which these are known oncogenic drivers using the Cancer Cell Line Encyclopedia (CCLE) mutations database (Broad Institute Cancer Dependency Map (DepMap) Project. Retrieved from DepMap Download Portal. May 2026 data release; (Ghandi et al., 2019)). Specifically, we looked at KRAS status in our CRC and PDAC panels, as well as mutations in the NFE2L2 (NRF2)-KEAP1 regulatory axis in our NSCLC LUAD and LUSC panels. For CRC we additionally annotated BRAF mutation status. Concerning KRAS/BRAF mutation status in the CRC panel (Fig. 3A), we see no clear pattern related to VitC sensitivity. Likewise, for the PDAC panel (Fig. 2A), while the cell lines display a range of varying VitC sensitivities, from CFPAC-1 with relatively low sensitivity to Mia Paca-2 with relatively high sensitivity, the large majority has a mutant KRAS background. For lung cancer we evaluated whether the metabolic liability caused by NRF2 activation and/or KEAP1 loss-of-function in these cell lines correlated with a heightened sensitivity to pro-oxidant VitC cell killing. While our LUSC panel (Fig. S3A) was too small (4 cell lines) to draw strong conclusions, we found no correlation for the LUAD panel (Fig. 5A), as both the least VitC sensitive (H2228) as well as 2 of the most VitC sensitive (H322 and A549) cell lines were characterized by an overactive NRF2 system (via NRF2 activating or KEAP1 loss-of-function mutations, respectively). These findings suggest that the genetic backgrounds of tumor cells have little influence on their respective susceptibility to VitC.

### 3.5. Pan-Cancer biology associated with VitC susceptibility

Our findings showed considerable heterogeneity at the level of individual proteins linked to VitC sensitivity per cancer cell panel, while strong common biology was shared between most of them. By studying the overlap between significantly enriched biological terms among cancer panels we found cell cycle, RNA metabolism, catalytic complex and biosynthesis to be associated with high VitC sensitivity (Fig. 6A). On the other hand, low VitC sensitivity was associated with extracellular vesicles/exosomes, mitochondrial terms, cell adhesion and protein transport terms across all panels (Fig. 6B).

**Fig. 6.**
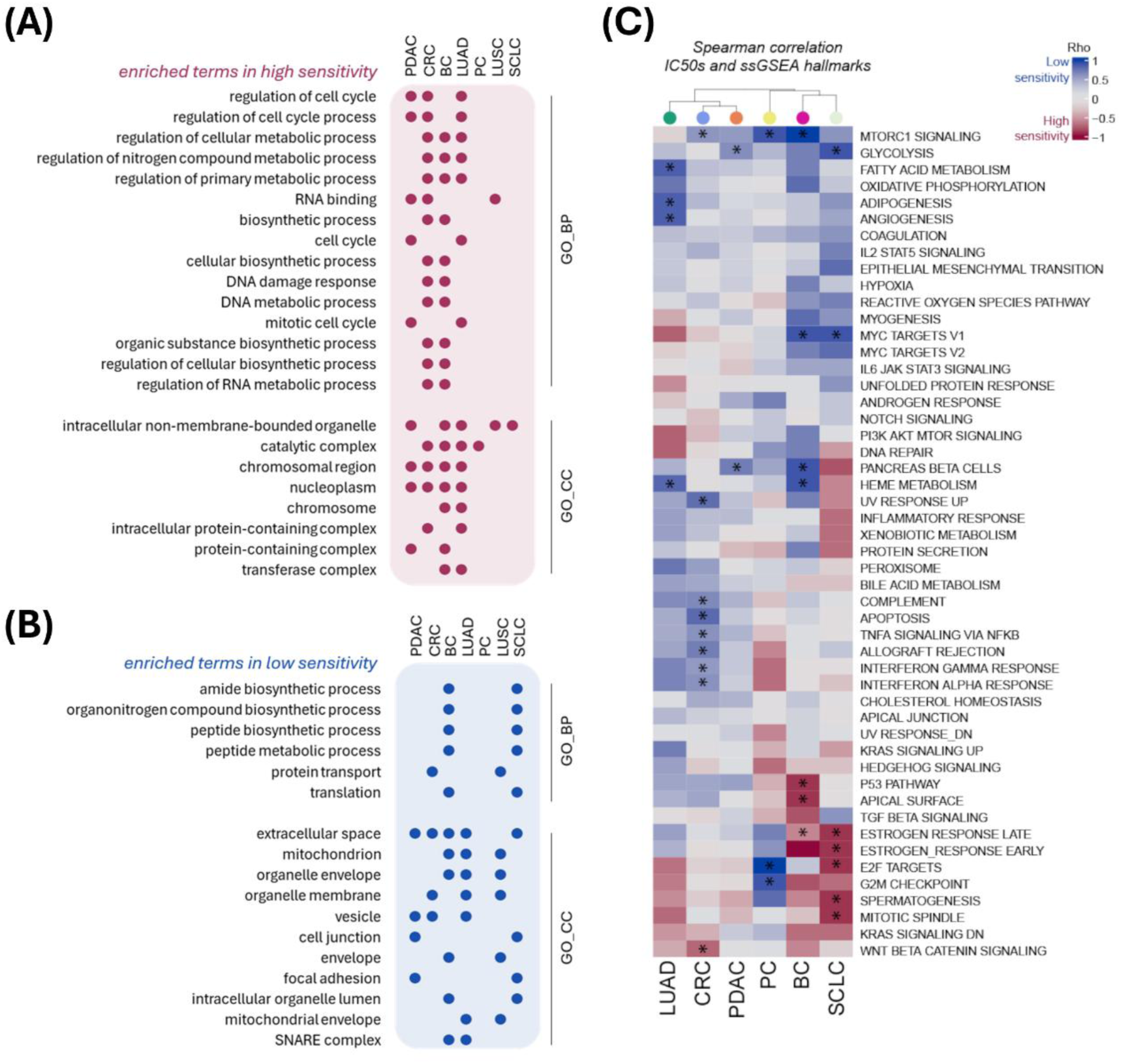
Enriched common biology associated with VitC sensitivity among cancer cell line panels. Overlapping significant (*pvalue adjusted*) gProfiler terms (GO:BP and GO:CC) in (A) highly sensitive cells and (B) less sensitive cells from shared proteins between spearman correlation and 2-group comparison analyses per panel. Maximum of top fifty terms ordered by adjusted *pvalue* were used to assess term overlap between panels. All shared terms were then summarized and redundancy was reduced using Revigo. (C) MSigDB hallmark signatures (ssGSEA) associated with VitC sensitivity across cancer panels. Heatmap built with spearman correlation coefficients (rho) between IC50s and ssGSEA hallmark enrichment scores per panel. Significant *pvalue* is annotated with an asterisk. LUSC panel was excluded from this analysis due to a low sample size (non-reliable results).

By interrogating the relationship of the enriched MSigDB hallmark signatures with VitC IC50s among panels, we observed 2 clusters of shared biology: adenocarcinomas PDAC, LUAD and CRC clustered away from PC, BC and SCLC (Fig. 6C). This difference could be influenced by hormone levels which are known to play a role in PC and BC, as well as in SCLC cell cultures (Sorenson et al., 1981). Overall, cancer cell lines with enriched proliferative hallmarks e.g. G2M checkpoint, mitotic spindle and Wnt beta catenin were more sensitive to VitC, whereas cell lines with strong survival signals like mTOR pathway, glycolysis, fatty acid metabolism were less sensitive, with Heme metabolism and Myc targets also enriched for a subset of panels. High proliferative signals were associated to high VitC sensitivity for all panels with the exception of the prostate cancer panel, which behaved in the opposite manner as observed earlier in the per panel analysis. Expression of proliferation marker MKI67 in more and less sensitive cell lines in different panels also showed this behaviour (Fig. S4A).

Merging all VitC associated proteins from the separate cancer cell panel analyses into one protein-protein interaction network, underscores the shared biology as can be evidenced by the high network connectivity (Fig. 7-8, Fig. S5-6). Pan-cancer network biology associated with high VitC sensitivity is shown in Figs.7 and S5 and low sensitivity in Figs. 8 and S6. These analyses confirm and extend the gene ontology and pathway enrichment analyses with more functional details.

**Fig. 7.**
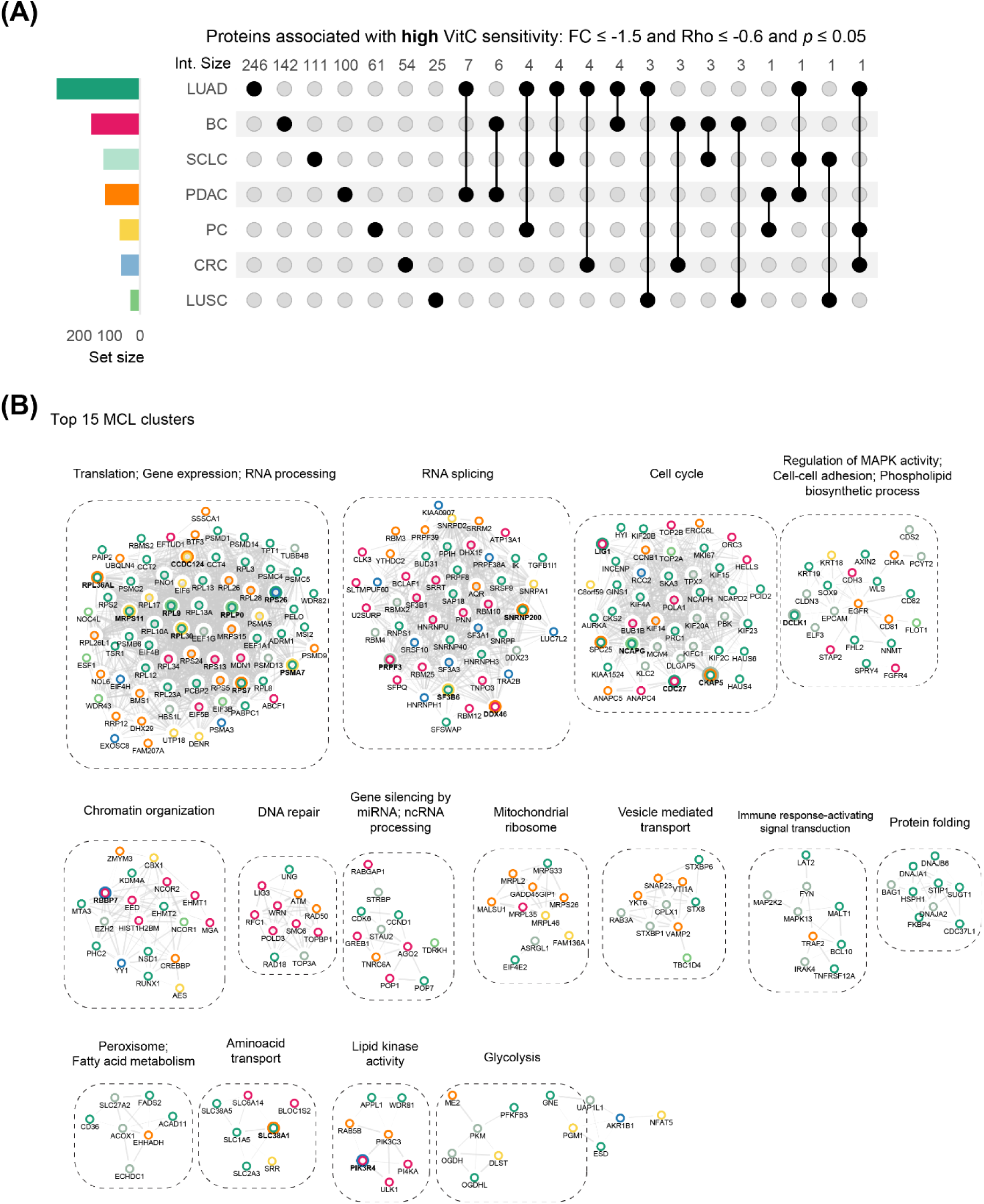
Pan-cancer biology associated with high VitC sensitivity. (A) Upset plot highlighting the overlap of proteins associated with high VitC sensitivity identified in each cancer panel. Overlap between panels is indicated with joined black circles. (B) Top 15 protein-protein interaction clusters and enriched biology. All proteins in panel A were used as input for the networks. Node colours refer to cell line panel, see Fig. 7A. Bold nodes refer to shared proteins between at least two cancer cell panels. See the expanded network in Fig. S5.

**Fig. 8.**
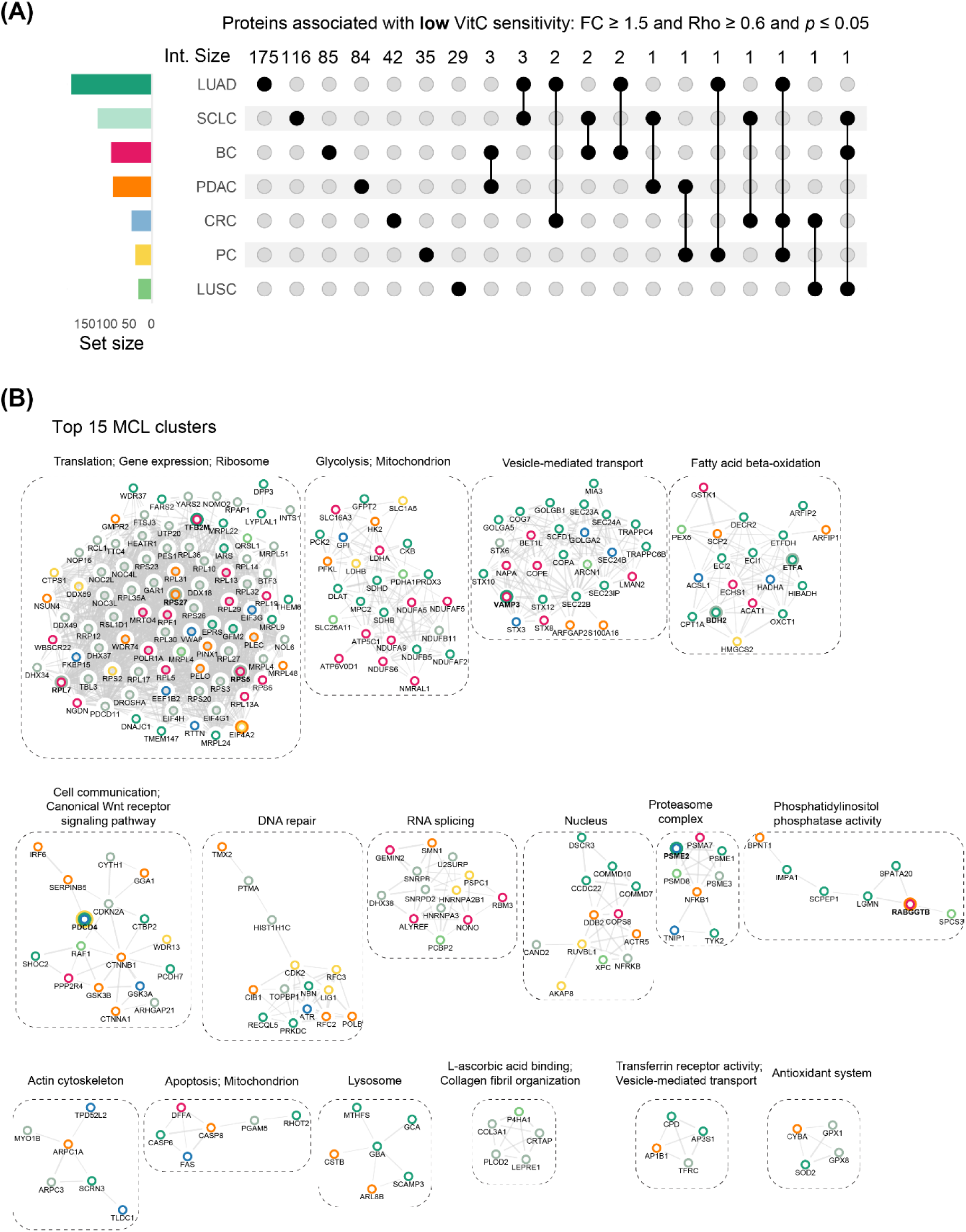
Pan-cancer biology associated with low VitC sensitivity. (A) Upset plot highlighting the overlap of proteins associated with low VitC sensitivity identified in each cancer panel. Overlap between panels is indicated with joined black circles. (B) Top 15 protein-protein interaction clusters and enriched biology. All proteins in panel A were used as input for the networks. Node colours refer to cell line panel, see Fig. 8A. Bold nodes refer to shared proteins between at least two cancer cell panels. See the expanded network in Fig. S6.

**Fig. 9.**
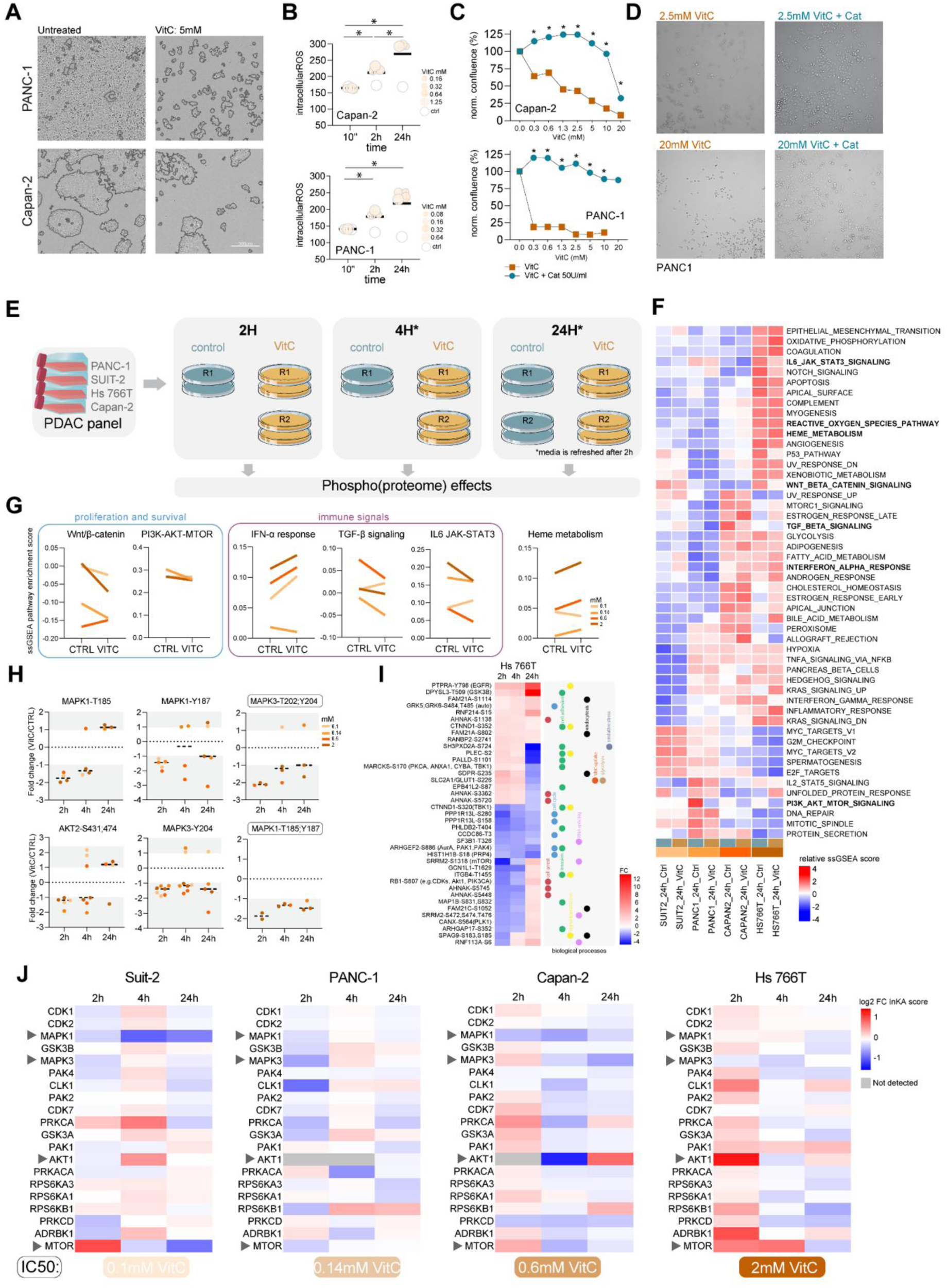
Effects of VitC on the proteome and phosphoproteome of a subset of PDAC cells. (A) Cell confluence images of PANC-1 and Capan-2 cell lines for treated (5mM) and untreated conditions. (B) Intracellular ROS levels as measured using OxiSelect™ Intracellular ROS Assay Kit upon multiple VitC concentrations in PANC-1 and Capan-2 cell lines after 10 minutes, 2h and 24h. For every time point, intracellular ROS levels were normalized to the levels of the untreated condition. Data from n=2 biological replicates. Statistical significance between timepoints was calculated by a paired t-test. (C) Cell growth of PANC-1 and Capan-2 upon VitC, and upon VitC followed by the exposure to catalase. (D) Representative images of PANC-1 cells VitC treated (2.5 and 20mM) with and without catalase exposure. (E) Workflow scheme of phosphoproteomic profiling of VitC treated cell lines at 2h, 4h and 24h. See Methods. (F) Unsupervised clustering of pathway enrichment scores generated by ssGSEA of the proteomics data, using MSigDB hallmark signatures. Enrichment scores are shown as z-scores. Processes marked in bold are shown as trend plots in subfigure (G). (G) Trend plots of selected ssGSEA enrichment pathways shown in (F) between control and VitC treatment groups after 24h. Colours depict cell lines in order of VitC sensitivity and VitC dose (0.1mM Suit-2, 0.14mM PANC-1, 0.6mM Capan-2, 2mM Hs 766T). (H) Phosphosite changes on MAPK1/3 and AKT2 upon VitC treatment at 2h, 4h and 24h timepoints, calculated by FC between control and VitC in all cell lines. Colours depict cell lines in order of VitC sensitivity and VitC dose (0.1mM Suit-2, 0.14mM PANC-1, 0.6mM Capan-2, 2mM Hs 766T). Rectangle indicates double phosphorylation required for MAPK1/3 activation. (I) Phosphosite changes upon VitC treatment at 2h, 4h and 24h timepoints, calculated by FC between control and VitC in Hs 766T cell line. Kinases responsible for the phosphorylation of certain target sites are indicated in brackets. (J) Kinase activity changes upon VitC treatment at 2h, 4h and 24h timepoints. Log2 fold changes were calculated for the top 20 ranked kinases based on the cumulative INKA score. A fold change cut-off of (-) 1.5 was used to determine which kinases are altered upon VitC treatment. In grey, FC values cannot be determined due to the absence of detected INKA kinases in at least one of the replicates, either in the VitC or control group. Arrows indicate key members of MAPK and PI3K/AKT signalling.

Overall, sensitive cancer cells exhibited higher expression of proteins involved in mRNA splicing and transcription, cell cycle, signal transduction and chromatin organization (Fig. 7). Another interesting protein cluster was involved in peroxisomal activity (ACAD11, ACOX1, SLC27A2, FADS2, EHHADH). Cancer cells with low sensitivity had elevated expression of proteins with functions in glycolysis and mitochondrial-related processes, vesicle-mediated transport, fatty-acid beta oxidation and Wnt receptor signalling pathway (Fig. 8). Importantly, we also identified two clusters with proteins involved in L-ascorbic acid binding and collagen fibril organization (PLOD2, COL3A1, P4HA1, LEPRE1) as well as Transferrin receptor activity linked to vesicle-mediated transport (TFRC, AP3S1, AP1B1, CPD) in the VitC low-sensitive group. Interestingly, a smaller vesicle-mediated transport cluster was found in sensitive cells, particularly expressed in the lung and pancreatic cancer panels. These proteins were involved in vesicle-membrane fusion (SNAP23, VTI1A, YKT6, STX8, STXBP1), exocytosis and secretion (RAB3A, STXBP6), as well as in insulin-dependent trafficking of GLUT4, which can be increased on the cell surface and enhance glucose transport (TBC1D4). Vesicle-mediated transport cluster identified in less sensitive cells showed a clearly distinct function, e.g. COPI (COPA, COPE), COPII (SEC24A, SEC24B, SEC23A, SEC23IP), and TRAPP (TRAPPC4, TRAPPC6B) complexes, and SNAP receptor activity and SNARE binding activity (NAPA, BET1L, STX12, STX6, STX10, VAMP3). Interestingly, retrograde golgi-to-ER transport proteins (ARCN1, STX8, SCFD1) and glycosylation proteins (LMAN2, GOLGA2) were also identified.

Among the proteins shared in more than two or three panels (Fig. 7A, Fig. 8A), two proteins, HCCS and PDCD4, were found to be significantly associated with low sensitivity in 3 cancer cell panels (Fig. S4B). Interestingly, HCCS is a lyase that catalyses the transfer of a mitochondrial heme group for the functioning of cytochrome c (Kranz et al., 2009). Given the importance of accessible free iron (Schoenfeld et al., 2017), we propose that elevated cytochrome c levels could suggest diminished levels of available iron, thereby reducing the pro-oxidant effect of high dose VitC. Besides this, decreased tumour suppressor PDCD4, found in 3 cancer cell panels with high VitC sensitivity (Fig. S4B), has been linked to increased proliferation in glioma cells (Pin et al., 2020), which supports our findings. In addition, other multiple-panel proteins associated with reduced VitC sensitivity were involved in vesicle-mediated transport (VAMP3), fatty acid oxidation (BDH2, ETFA), ERK signalling (RPS6KA3), proteasome (PSME2), calcium homeostasis (S100A6) and mitochondrial homeostasis and respiration (HSPA9, COQ6) (Fig. 8, Fig. S6, see protein names highlighted in bold). Two pan-cancer proteins associated with high VitC sensitivity were the microtubule dynamics and mitosis regulator CKAP5 (Ali et al., 2023; Miller et al., 2016), and MORF4L2, implicated in chromatin assembly and histone modification (Fig. S4B). Other proteins shared between sensitive cell lines from multiple cancer panels were also found to be involved in cell cycle (NCAPG, LIG1, PDS5A, TMPO, PPP1CB, PPME1, CAPRIN1, PKP4, SYNE2), RNA splicing (SF3B6, DDX46, SNRNP200, HCFC1, THOC5, PRPF38B) and chromatin organization (RBBP7, HCFC1) (Fig. 7, Fig. S5).

### 3.6. (Phospho-)proteomic changes upon high-dose VitC treatment

To assess the effect of high dose VitC on the proteome and phosphoproteome in a tumour type with poor prognosis, a subset of the PDAC panel, displaying a broad range of IC50 values, was analysed in more detail. Given the poor prognosis of this tumour type, we aimed to further elucidate the mechanism of action by examining on-treatment samples, thereby gaining a deeper understanding on tumour cell responses to VitC. First, we confirmed an increase in intracellular ROS levels with time on treatment (Fig. 9A-B), and a complete recovery of cell growth was achieved upon the addition of catalase, a well-known hydrogen peroxide scavenger (Fig. 9C-D). Subsequently, we performed phosphoproteomic profiling of four PDAC cell lines treated with VitC at their respective IC50 values at three distinct time points (2 hours, 4 hours, and 24 hours, Fig. 9E). This approach allowed us to capture early changes in the phosphoproteome as well as expression changes in the proteome. From the 38 samples, 3 samples were excluded from further analyses due to the low amount of phosphosites measured compared to replicates and external control sample HCT116 (Fig. S7A,B). Thus, a total of 35 samples, with an average of 3296 proteins and 7617 phosphosites per sample (Fig. S7C) were subsequently used in unsupervised clustering (Fig. S7D,E). By performing an unsupervised clustering, we observed that data primarily clustered based on the cell line type, and to a lower extent by treatment time point (Fig.S7D,E). Therefore, we followed a cell line-specific approach for data analysis. This approach allowed us to identify and compare cell line-specific mechanisms in all cell lines in order to uncover both unique and common changes in the (phospho-)proteome induced by VitC treatment.

Looking at the global protein expression data after 24h (Fig. 9F,G), we found a general decrease in proliferation and survival pathways such as Wnt/ß-catenin and PI3K-AKT-MTOR upon high-dose VitC treatment. Immune-related IFN-α response on the other hand was enhanced upon VitC treatment for 3 of the 4 cells lines tested. Interestingly, IFN-α response can be activated in tumour cells upon cell damage, which can trigger activation of macrophages, NK cells, and T cells, and eventually promotes cancer cell killing. This is in line with *in vivo* studies reporting increased IFN-α upon VitC anti-viral immune responses (Kim et al., 2013), as well as increased IFN-γ upon the synergistic combination of VitC coupled to immune checkpoint blockade (Magrì et al., 2020). Similarly, upon 24h of VitC treatment, the majority of cell lines showed a decrease in TGF-β and IL6 JAK-STAT3 signalling, two key immunosuppressive pathways in PDAC tumours that can positively regulate and attract myeloid-derived suppressor cells and regulatory T cells to the tumour microenvironment (Ho et al., 2020). Furthermore, as mentioned, Wnt/β-catenin signalling, which also holds an immunosuppressive role in PDAC tumours (Du et al., 2023), was found to be reduced upon VitC, in line with previous studies in PDAC (Kim et al., 2022).

By interrogating individual proteins, we observed only a few significant changes in the proteome after 24h. This suggests that a longer treatment duration (∼48h) may be necessary to capture significant alterations in the proteome. Despite this, we were able to identify interesting regulations, such as the decrease of iron receptor TFRC and IMMT expression (maintains the mitochondrial cristae morphology), or the increase of CYCS expression, which may be an indicator of mitochondrial outer membrane permeabilization (Garrido et al., 2006) (Fig. S7F). Overall, proteome changes suggested that after 24h high-dose VitC treatment, the majority of PDAC cell lines displayed reduced immunosuppressive and survival signals, and overall increased iron metabolism, in line with reported mechanisms in multiple cancer types (Böttger et al., 2021).

Concerning the phosphoproteomics data, most identified phosphosite changes occurred on the cell line treated with the highest VitC IC50 dose, i.e. Hs 766T with 2mM (Fig. 9I). In particular, we observed a significant regulatory impact on various phosphorylation events on proteins involved in cell cycle arrest (RB1, AHNAK), cell cycle progression (e.g. GRK5, CCDC86), cell adhesion and invasion (e.g. CTNND1, PALLD, ITGB4), signal transduction (e.g. PTPRA, SPAG9) and endocytosis (e.g. FAM21, SDPR). Phosphorylation site changes of particular significance were identified in proteins including GLUT1/SLC2A1, RB1, MARCKS, PTPRA, and CTNND1. Specifically, the phosphorylation site S266 of GLUT1 exhibited an upregulation at 2 hours and 4 hours, followed by a downregulation at 24 hours. Notably, this phosphorylation event is known to enhance the translocation of GLUT1 to the plasma membrane, facilitating glucose uptake (Lee et al., 2014). The fact that GLUT1 phosphorylation was upregulated in the early hours of treatment may suggest that oxidized VitC (DHA) and glucose are competing for cell uptake. Thus, its subsequent downregulation at 24h could be a cell-intrinsic mechanism to reduce the VitC uptake. Likewise, phosphorylation of FAM21A S802/S1114 and FAM21C S1052, both proteins involved in recycling GLUT1 (Lee et al., 2016), were found to be upregulated over time. This upregulation holds significance, considering that GLUT1 is essential for transporting DHA into the cells.

For the cells treated with lower doses, we also observed a general increase in phosphorylation occurring in cell cycle arrest proteins after 24h (e.g. AHNAK), as well as in cell adhesion markers (e.g. LMNA, FLNB, FMNL2) and splicing regulators (e.g. SRRM2) (data not shown). Interestingly, phosphorylation on mTORC2 subunit RICTOR-T1332, was downregulated in Capan-2, and negative mTOR-regulatory AKT1S1-S203 was upregulated in Suit-2.

By looking at relevant downstream regulators of KRAS mutant tumours, e.g. MAPK and PI3K/AKT pathways, we observed a broad downregulation upon treatment, most notably at early time points (Fig. 9H; negative fold change compared to control upon treatment). The intensities of double phosphorylations on MAPK3-T202/Y204 and MAPK1-T185/Y187, which are important for ERK pathway activation, as well as single phosphorylations at MAPK1-Y187, MAPK1-Y185 and MAPK3-Y204, were down-regulated at 2h at all VitC doses tested. Importantly, the observed average phosphosite changes for MAPK1 and 3 across all 4 PDAC cell lines were also reflected in MAPK1/3 activity per cell line (Fig. 9J). When kinase activity changes were evaluated upon VitC treatment over time using the InKA pipeline, MAPK1 and/or MAPK3 kinase activity showed a consistent decrease across all time points, most notably in Suit-2 and Capan-2.

AKT1 phosphosites were not identified in our set up, however we detected AKT2 S431/474, which showed decreased phosphorylation at 2 and 4 hours for PANC-1, while Capan-2 and Hs 766T remained nearly unchanged (Fig. 9H). The cell line most sensitive to VitC, Suit-2, on the contrary, showed an increase 4 hours after treatment. Interestingly, AKT2-S474 is known to be crucial for insulin-stimulated mTOR activation and glucose uptake in adipocytes (Kearney et al., 2019). While AKT1 kinase activity changes could not be determined for PANC-1 and Capan-2 at 2h (Fig. 9J), a downregulation was observed in Capan-2 at 4 hours, followed by an upregulation at 24 hours. For Hs 766T, a slight down-regulation of AKT1 at 4h returned to control levels after 24h. The most consistent change was mTOR kinase activity (Fig. 9J), which declined over time in most cell lines, consistent with the reduced activity of the PI3K-AKT-MTOR pathway at the protein level (Fig. 9G). These findings are in line with previous studies in thyroid (Su et al., 2019) and breast (Mustafi et al., 2021) cancer cells showing inhibition of PI3K-AKT-MTOR and MAPK pathways upon high-dose VitC, further supporting our results.

Although not in the top 20 rankings, kinase activity of CAMK2D and CAMK2A, involved in antioxidant mechanisms by sensing ROS-meditated stress levels (Erickson et al., 2011), were downregulated upon VitC treatment, particularly after 24h, in both Capan-2 and PANC-1 cell lines (data not shown). This data may suggest that antioxidant mechanisms are overcome after 24h, in line with pro-oxidant-mediated cytotoxicity (Fig. 9A,B).

## 4. DISCUSSION

In high concentrations (mM range), which can only be reached by intravenous administration, VitC has proved to exert potent anti-cancer effects in multiple cancer types. Here we report VitC IC50 values in relation to proteome features and associated biological functions for a large pan-cancer cell panel (n=51) representing major solid cancer types.

Importantly, all cancer cell lines tested in this study were particularly sensitive, with an average IC50 of 1.7±0.4 mM, with only 5 out of 51 cell lines requiring a 5-10 mM dose. Overall, all tested IC50 values were within the clinically achievable range, in line with previous studies (Böttger et al., 2021; Chen et al., 2008; Schoenfeld et al., 2017). Furthermore, our results for PDAC, CRC, BC and LC also showed that VitC sensitivity is independent of cancer subtypes (e.g. epithelial/mesenchymal or CMS subtypes) or genetic aberrations, which positions VitC as a promising one-size-fits-all anti-cancer agent. The baseline proteome data showed that high VitC sensitivity is associated with proliferation-related functions, while lower sensitivity is linked to metabolic processes and vesicle-mediated transport. Finally, (phospho-)proteome analysis of on-treatment VitC effects revealed down-modulation of MAPK1/2, AKT-MTOR signalling and immune suppressive signalling, while IFN-α response was enhanced upon VitC.

### 4.1. Confirmation of well-established VitC effects

By exploring known proteins associated with VitC susceptibility in cancer, we were able to confirm previous insights on the importance of iron metabolism, antioxidant system, ROS-dependent pathways, metabolic processes and DNA damage proteins. In particular, high doses of VitC are known to utilize intracellular iron levels to increase ROS levels. As a result, cancer cells with elevated free labile iron levels and low catalase are generally more susceptible to the pro-oxidant VitC cytotoxic effect. By interrogating known proteins involved in iron metabolism, we found iron transport members like DNMT1, ZIP14 and FLVCR1 were found to be increased in more sensitive cell lines, which underscores the role of iron trafficking and the intracellular free iron pool as the key player in VitC sensitivity. These members are involved in non-TF-bound iron pathway, typically occurring during iron overload (Xu et al., 2012). Remarkably, iron importer TFRC was found to be enriched in less sensitive cell lines. We hypothesize that balanced expression of these members is crucial for increased labile iron pool levels, which are crucial to promoting the VitC pro-oxidant effect in cancer cells, as previously described (Schoenfeld et al., 2017).

### 4.2. Proliferative functions

Findings from both individual cancer panels and our comprehensive pan-cancer panel analyses indicate an association between proliferative functions and high sensitivity to VitC. Notably, proliferative signals like elevated EGFR and IGF2R were also linked to increased VitC sensitivity in the PDAC panel. In line with this, we identified four potential pan-cancer proteins highly associated with VitC sensitivity, with mitotic, chromatin and histone regulators CKAP5 and MORF4L2 specifically found to be increased in highly sensitive cell lines in SCLC, LUAD and PDAC, and LUAD, CRC and PC panels, respectively. Interestingly, a pan-cancer protein found to be reduced in sensitive cell lines in BC, LUSC and SCLC panels was PDCD4, a tumour suppressor found to be reduced in highly proliferative glioma cells (Pin et al., 2020), consistent with our findings.

### 4.3. Mitochondrial functions, iron and oxidative stress levels

Cancer cells with VitC IC50 values in the mM range exhibited, as expected, higher abundance of antioxidant defence proteins, as well as higher levels of proteins with vesicle transport and mitochondrial functions. For instance, SPR protein was found to be associated with low VitC sensitivity in the CRC panel. This protein is a promoter of BH4, which is a potent antioxidant that may be involved in ferroptosis induction (Soula et al., 2020). Intriguingly, BH4 levels and its antioxidant activity can regulate iron metabolism and mitochondrial function of T cells too. From the pan-cancer network analysis, multiple antioxidants such as CYBA, GPX1, SOD2 and GPX8 were also identified in less sensitive cell lines. In the BC panel, GSTK1, which is an antioxidant contained in peroxisomes, correlated with low VitC sensitivity, and has been found to be involved in ferroptosis together with CAT, SOD1 and PRDX5 (Chen et al., 2021). This cancer cell death process has been linked to VitC anti-cancer effect in several recent studies and may be further studied (Liu et al., 2022; Wang et al., 2021).

These metabolic and mitochondrial functions were confirmed in the common pan-cancer analysis, which supports that energy production processes can also be targeted by VitC (El Hassouni et al., 2020). Wnt beta catenin signalling significantly correlated with high VitC sensitivity in the CRC panel, however from the network analysis we also identified members of this pathway (e.g. CTNNA1, CTNNB1, GSK3A/B) correlating with low sensitivity in other panels. To some extent, beta catenin could be involved in VitC sensitivity in cancer cells since it plays a key role in heme synthesis, glucose uptake and fatty acid metabolism, and positively regulates L-gulonolactone oxidase, crucial for VitC biosynthesis (Goel et al., 2022). Furthermore, VitC has been recently shown to inhibit EMT in PDAC via the Wnt/beta catenin pathway (Kim et al., 2022), which underscores VitC’s multi-targeting capability in cancer. Besides, mTOR was the pathway most significantly associated with reduced VitC sensitivity. This pathway is key for cancer survival, and it has recently been shown to be inhibited by VitC *in vivo* (Qin et al., 2023).

A pan-cancer protein associated with lower VitC sensitivity was HCCS, which catalyses the heme group linking, crucial for cytochrome c synthesis. We suggest that mature cytochrome c has the ability to associate with free iron, consequently mitigating oxidative stress induced by high doses of VitC. High-dose VitC is known to reduce cytochrome c levels, leading to impaired mitochondrial respiration, reducing ATP levels and yielding superoxide (Bakalova et al., 2020). However, cancer cells are able to surpass mitochondrial ATP production by promoting aerobic glycolysis, known as the Warburg effect. Our findings reveal an enrichment of this process in cancer cells exhibiting diminished sensitivity to VitC. These mechanisms, targeting the energy levels and increasing reactive oxygen species, together with sensitivity associated proliferative signals, make high-dose VitC a very promising candidate for anti-cancer therapy comparable to chemotherapeutics or mitochondrial targeting agents, yet with reduced side effects.

Furthermore, our findings position VitC as a potential treatment for chemoresistant tumours. In the LUAD panel, the cell line resistant to cisplatin was highly sensitive to VitC. Interestingly, PDAC’s top candidate associated with higher VitC sensitivity was ANPEP, a protein associated with multi-drug resistance, also to chemotherapeutic drugs, in multiple cancer types (Pizzo et al., 2023). Other proteins involved in chemoresistance, namely TGM2 and VCP, were associated with high VitC sensitivity in the LUAD panels. When considering our evaluation of cisplatin responses across the LUAD cell line panel, this further underscores VitC as a potential alternative treatment for chemoresistant tumours.

Altogether, these findings position high-dose VitC as a broad anti-cancer agent comparable to chemotherapeutic or mitochondrial disruptor agents. Particularly for heavily treated patients with hard-to-treat tumours such as PDAC, CRC or NSCLC, a vast amount of preclinical evidence supports high-dose intravenous VitC as a promising alternative (Böttger et al., 2021). Promising phase I/II clinical studies are continuously showing VitC as a tolerable, safe and effective anti-cancer agent in combination with standard treatments, e.g. in NSCLC and GBM (Schoenfeld et al., 2017) and in PDAC (Welsh et al., 2013). Larger scale randomized phase III clinical studies are warranted to support its clinical efficacy.

### 4.4. On-treatment VitC effects in PDAC cell lines

High dose VitC at IC50 values induced both cell-specific alterations as well as common down-modulation of cell survival signalling and differential effects on immune signalling, shifting cells to less immune suppression. In contrast to the high number of phosphosite alterations after 2-4h, we noted only a limited number of changes in protein expression following 24 hours of VitC treatment. The relatively brief duration for detecting protein expression changes, coupled with the use of lower doses (< 1 mM), may explain why we did not see changes in EMT hallmark proteins, e-cadherin or vimentin, contrary to what was previously described in PDAC cell lines (Polireddy et al., 2017). Possibly, more protein expression changes may be found by including later timepoints (>48h), in particular those related to tumour survival and growth.

### 4.5. Limitations of the study

This study is of a descriptive nature, and our large scale pharmacological and proteomics datasets should be seen as a resource for further research. Although our results highlight pathways represented in more or less VitC-sensitive cell lines, there is little mechanistic information about their relevance in determining sensitivity to VitC. Heading forward, perturbation experiments with knock-downs or knock-outs would have to be performed to see if we can further sensitize cancer cell lines that are less sensitive and thereby test the premise that proteins and pathways that are enriched in more VitC-sensitive cell lines point to mechanisms of sensitivity, while those enriched in less VitC-sensitive lines underlie mechanisms of resistance. Of note, we do not call these less sensitive cell lines “resistant” as cell growth can be inhibited at a clinically achievable dose in all tested cell lines.

Furthermore, as this study involved a large panel of cell lines representing multiple cancer types, concessions had to be made concerning the culturing conditions. We decided against a single compromise medium for all cell lines in order to avoid phenotypic drift and suboptimal viability. The limitation of this approach is that different media formulations contain distinct concentrations of nutrients, growth factors, and metabolites, which introduces confounding variables when testing VitC sensitivities. Specifically, the rate of ascorbate oxidation can vary between the two different base medium formulations (DMEM and RPMI 1640) used, primarily due to differences in iron, riboflavin and glutathione concentrations, leading to differences in hydrogen peroxide (H₂O₂) and reactive oxygen species (ROS) generation. DMEM inherently exhibits a higher basal redox activity compared to RPMI 1640 (Zwolak et al., 2022). Importantly, while the original iron distinction between the 2 formulations becomes significantly masked and biologically diluted upon the addition of serum (FCS), other factors such as DMEM’s higher concentration of riboflavin and RPMI’s built-in glutathione buffer still mean that DMEM has the potential to generate artificial peaks of H₂O₂ faster than RPMI. We have tried to address this caveat by applying only a brief VitC exposure window of 2h (refreshing the medium thereafter) in our cell viability assays, thus limiting the extracellular generation of H₂O₂ and other byproducts in the presence of trace metals.

## 5. CONCLUSION

Our findings showed remarkable VitC sensitivity of all cell lines within the clinically achievable range, suggesting that predictive markers are unnecessary. The pro-oxidant effects and high proliferation signature observed in highly sensitive cell lines provides valuable support for positioning VitC as an alternative agent to chemotherapy, which is also shown to specifically target cancer cells rather than normal cells (Chen et al., 2008; Schoenfeld et al., 2017). Importantly, our findings indicate that sensitivity to VitC and sensitivity to other cytotoxic agents is not one-to-one correlated, and that VitC toxicity is not dependent on underlying histological or genetic subtypes of cancers. Moreover, the significant increase in antioxidant and mitochondrial activity observed in less sensitive cell lines reinforces the well-established mechanism of action of VitC. Finally, the on-treatment analysis in a subset of the PDAC panel, revealed changes in the phosphoproteome and kinase activity involving proliferation and survival, invasion and immune signals. The findings from this study underscore VitC’s effectiveness against multiple cancer types, independent of subtypes, thereby laying the foundation for positioning this compound as a promising anti-cancer agent that warrants wide-spread clinical studies, as well as use in the palliative setting.

## ACKNOWLEDGEMENTS

Maarten Bijlsma is acknowledged for providing pancreatic cancer cell lines. Michiel Pegtel and Caitrin Crudden are acknowledged for providing breast cancer cell lines. Cancer Center Amsterdam and Netherlands Organisation for Scientific Research (NWO-Middelgroot project number 91116017) are acknowledged for support of the mass spectrometry infrastructure and Surfsara for computing infrastructure (reference e-infra180166).

## AUTHOR CONTRIBUTIONS

A.V.M. and F.B. designed the study, performed experiments, analysed data, performed bioinformatic analyses, and wrote the manuscript. K.T., E.H.Y.Y, L.M., S.S. and D.D. were involved in cell culture and VitC sensitivity experiments. M. M., T.L.L., I.B., A.E. and T.S. were involved in proteomic sample processing. S.R.P. performed the mass spectrometry measurements and S.R.P., T.V.P. and J.C.K. analysed data. E.G. and J.P.M. collaborated by providing cell lines. C.R.J. designed and supervised the study and wrote the manuscript.

## PEER REVIEW

## DATA ACCESSIBILITY

The mass spectrometry proteomics data have been deposited to the ProteomeXchange Consortium via the PRIDE (Perez-Riverol et al., 2022) partner repository with the dataset identifier PXD046089. Human Swiss-Prot database used for raw data search was acquired from the UniProt database (https://www.uniprot.org/).

## ABBREVIATIONS

VitC: vitamin C
PDAC: pancreatic ductal adenocarcinoma
CRC: colorectal cancer
LUAD: lung adenocarcinoma
LUSC: lung squamous cell carcinoma
SCLC: small cell lung cancer
PC: prostate cancer
BC: breast cancer
ROS: reactive oxygen species

## REFERENCES

Ali A, Vineethakumari C, Lacasa C, Lüders J. 2023. Microtubule nucleation and γTuRC centrosome localization in interphase cells require ch-TOG. Nature Communications 2023 14:1 14:1–17. DOI: 10.1038/s41467-023-35955-w, PMID: 36702836

Ashburner M, Ball CA, Blake JA, Botstein D, Butler H, Cherry JM, Davis AP, Dolinski K, Dwight SS, Eppig JT, Harris MA, Hill DP, Issel-Tarver L, Kasarskis A, Lewis S, Matese JC, Richardson JE, Ringwald M, Rubin GM, Sherlock G. 2000. Gene Ontology: tool for the unification of biology. Nature Genetics 2000 25:1 25:25–29. DOI: 10.1038/75556, PMID: 10802651

Aumailley L, Bourassa S, Gotti C, Droit A, Lebel M. 2021. Vitamin C Differentially Impacts the Serum Proteome Profile in Female and Male Mice. Journal of Proteome Research 20:5036–5053. DOI: 10.1021/ACS.JPROTEOME.1C00542/ASSET/IMAGES/LARGE/PR1C00542_0008.JPEG, PMID: 34643398

Bakalova R, Zhelev Z, Miller T, Aoki I, Higashi T. 2020. Vitamin C versus Cancer: Ascorbic Acid Radical and Impairment of Mitochondrial Respiration? Oxidative Medicine and Cellular Longevity 2020. DOI: 10.1155/2020/1504048, PMID: 32411317

Beekhof R, Alphen C van, Henneman AA, Knol JC, Pham T V, Rolfs F, Labots M, Henneberry E, Large TY Le, Haas RR de, Piersma SR, Vurchio V, Bertotti A, Trusolino L, Verheul HM, Jimenez CR. 2019. INKA, an integrative data analysis pipeline for phosphoproteomic inference of active kinases. Molecular Systems Biology 15:e8250. DOI: 10.15252/MSB.20188250, PMID: 30979792

Böttger F, Schaaij-Visser TB, de Reus I, Piersma SR, Pham T V., Nagel R, Brakenhoff RH, Thunnissen E, Smit EF, Jimenez CR. 2019. Proteome analysis of non-small cell lung cancer cell line secretomes and patient sputum reveals biofluid biomarker candidates for cisplatin response prediction. Journal of Proteomics 196:106–119. DOI: 10.1016/J.JPROT.2019.01.018, PMID: 30710758

Böttger F, Vallés-Martí A, Cahn L, Jimenez CR. 2021. High-dose intravenous vitamin C, a promising multi-targeting agent in the treatment of cancer. Journal of Experimental & Clinical Cancer Research : CR 40:343. DOI: 10.1186/S13046-021-02134-Y, PMID: 34717701

Brown RAM, Richardson KL, Kabir TD, Trinder D, Ganss R, Leedman PJ. 2020. Altered Iron Metabolism and Impact in Cancer Biology, Metastasis, and Immunology. Frontiers in Oncology 10:533570. DOI: 10.3389/FONC.2020.00476/BIBTEX

Buranasudja V, Doskey CM, Gibson AR, Wagner BA, Du J, Gordon DJ, Koppenhafer SL, Cullen JJ, Buettner GR. 2019. Pharmacologic ascorbate primes pancreatic cancer cells for death by rewiring cellular energetics and inducing DNA damage. Molecular Cancer Research 17:2102–2114. DOI: 10.1158/1541-7786.MCR-19-0381/82107/AM/PHARMACOLOGICAL-ASCORBATE-PRIMES-PANCREATIC-CANCER, PMID: 31337671

Chen Q, Espey MG, Krishna MC, Mitchell JB, Corpe CP, Buettner GR, Shacter E, Levine M. 2005. Pharmacologic ascorbic acid concentrations selectively kill cancer cells: Action as a pro-drug to deliver hydrogen peroxide to tissues. Proceedings of the National Academy of Sciences of the United States of America 102:13604 LP – 13609.

Chen Q, Espey MG, Sun AY, Pooput C, Kirk KL, Krishna MC, Khosh DB, Drisko J, Levine M. 2008. Pharmacologic doses of ascorbate act as a prooxidant and decrease growth of aggressive tumor xenografts in mice. Proceedings of the National Academy of Sciences 105:11105–11109. DOI: 10.1073/pnas.0804226105, PMID: 18678913

Chen X, Kang R, Kroemer G, Tang D. 2021. Organelle-specific regulation of ferroptosis. Cell Death & Differentiation 2021 28:10 28:2843–2856. DOI: 10.1038/s41418-021-00859-z, PMID: 34465893

Consortium TGO, Aleksander SA, Balhoff J, Carbon S, Cherry JM, Drabkin HJ, Ebert D, Feuermann M, Gaudet P, Harris NL, Hill DP, Lee R, Mi H, Moxon S, Mungall CJ, Muruganugan A, Mushayahama T, Sternberg PW, Thomas PD, Van Auken K, Ramsey J, Siegele DA, Chisholm RL, Fey P, Aspromonte MC, Nugnes MV, Quaglia F, Tosatto S, Giglio M, Nadendla S, Antonazzo G, Attrill H, dos Santos G, Marygold S, Strelets V, Tabone CJ, Thurmond J, Zhou P, Ahmed SH, Asanitthong P, Luna Buitrago D, Erdol MN, Gage MC, Ali Kadhum M, Li KYC, Long M, Michalak A, Pesala A, Pritazahra A, Saverimuttu SCC, Su R, Thurlow KE, Lovering RC, Logie C, Oliferenko S, Blake J, Christie K, Corbani L, Dolan ME, Drabkin HJ, Hill DP, Ni L, Sitnikov D, Smith C, Cuzick A, Seager J, Cooper L, Elser J, Jaiswal P, Gupta P, Jaiswal P, Naithani S, Lera-Ramirez M, Rutherford K, Wood V, De Pons JL, Dwinell MR, Hayman GT, Kaldunski ML, Kwitek AE, Laulederkind SJF, Tutaj MA, Vedi M, Wang S-J, D’Eustachio P, Aimo L, Axelsen K, Bridge A, Hyka-Nouspikel N, Morgat A, Aleksander SA, Cherry JM, Engel SR, Karra K, Miyasato SR, Nash RS, Skrzypek MS, Weng S, Wong ED, Bakker E, Berardini TZ, Reiser L, Auchincloss A, Axelsen K, Argoud-Puy G, Blatter M-C, Boutet E, Breuza L, Bridge A, Casals-Casas C, Coudert E, Estreicher A, Livia Famiglietti M, Feuermann M, Gos A, Gruaz-Gumowski N, Hulo C, Hyka-Nouspikel N, Jungo F, Le Mercier P, Lieberherr D, Masson P, Morgat A, Pedruzzi I, Pourcel L, Poux S, Rivoire C, Sundaram S, Bateman A, Bowler-Barnett E, Bye-A-Jee H, Denny P, Ignatchenko A, Ishtiaq R, Lock A, Lussi Y, Magrane M, Martin MJ, Orchard S, Raposo P, Speretta E, Tyagi N, Warner K, Zaru R, Diehl AD, Lee R, Chan J, Diamantakis S, Raciti D, Zarowiecki M, Fisher M, James-Zorn C, Ponferrada V, Zorn A, Ramachandran S, Ruzicka L, Westerfield M, Aleksander SA, Balhoff J, Carbon S, Cherry JM, Drabkin HJ, Ebert D, Feuermann M, Gaudet P, Harris NL, Hill DP, Lee R, Mi H, Moxon S, Mungall CJ, Muruganugan A, Mushayahama T, Sternberg PW, Thomas PD, Van Auken K, Ramsey J, Siegele DA, Chisholm RL, Fey P, Aspromonte MC, Nugnes MV, Quaglia F, Tosatto S, Giglio M, Nadendla S, Antonazzo G, Attrill H, dos Santos G, Marygold S, Strelets V, Tabone CJ, Thurmond J, Zhou P, Ahmed SH, Asanitthong P, Luna Buitrago D, Erdol MN, Gage MC, Ali Kadhum M, Li KYC, Long M, Michalak A, Pesala A, Pritazahra A, Saverimuttu SCC, Su R, Thurlow KE, Lovering RC, Logie C, Oliferenko S, Blake J, Christie K, Corbani L, Dolan ME, Drabkin HJ, Hill DP, Ni L, Sitnikov D, Smith C, Cuzick A, Seager J, Cooper L, Elser J, Jaiswal P, Gupta P, Jaiswal P, Naithani S, Lera-Ramirez M, Rutherford K, Wood V, De Pons JL, Dwinell MR, Hayman GT, Kaldunski ML, Kwitek AE, Laulederkind SJF, Tutaj MA, Vedi M, Wang S-J, D’Eustachio P, Aimo L, Axelsen K, Bridge A, Hyka-Nouspikel N, Morgat A, Aleksander SA, Cherry JM, Engel SR, Karra K, Miyasato SR, Nash RS, Skrzypek MS, Weng S, Wong ED, Bakker E, Berardini TZ, Reiser L, Auchincloss A, Axelsen K, Argoud-Puy G, Blatter M-C, Boutet E, Breuza L, Bridge A, Casals-Casas C, Coudert E, Estreicher A, Livia Famiglietti M, Feuermann M, Gos A, Gruaz-Gumowski N, Hulo C, Hyka-Nouspikel N, Jungo F, Le Mercier P, Lieberherr D, Masson P, Morgat A, Pedruzzi I, Pourcel L, Poux S, Rivoire C, Sundaram S, Bateman A, Bowler-Barnett E, Bye-A-Jee H, Denny P, Ignatchenko A, Ishtiaq R, Lock A, Lussi Y, Magrane M, Martin MJ, Orchard S, Raposo P, Speretta E, Tyagi N, Warner K, Zaru R, Diehl AD, Lee R, Chan J, Diamantakis S, Raciti D, Zarowiecki M, Fisher M, James-Zorn C, Ponferrada V, Zorn A, Ramachandran S, Ruzicka L, Westerfield M. 2023. The Gene Ontology knowledgebase in 2023. Genetics 224. DOI: 10.1093/GENETICS/IYAD031

Cortés ÁA, Urquizu LC, Cubero JH. 2015. Adjuvant chemotherapy in non-small cell lung cancer: state-of-the-art. Translational lung cancer research 4:191–197. DOI: 10.3978/J.ISSN.2218-6751.2014.06.01, PMID: 25870801

Costantini S, Capone F, Polo A, Bagnara P, Budillon A. 2021. Valosin-Containing Protein (VCP)/p97: A Prognostic Biomarker and Therapeutic Target in Cancer. International journal of molecular sciences 22. DOI: 10.3390/IJMS221810177, PMID: 34576340

Cox J, Mann M. 2008. MaxQuant enables high peptide identification rates, individualized p.p.b.-range mass accuracies and proteome-wide protein quantification. Nature Biotechnology 26:1367–1372. DOI: 10.1038/nbt.1511, PMID: 19029910

Dai X, Cheng H, Bai Z, Li J. 2017. Breast Cancer Cell Line Classification and Its Relevance with Breast Tumor Subtyping. Journal of Cancer 8:3131. DOI: 10.7150/JCA.18457, PMID: 29158785

Doskey CM, Buranasudja V, Wagner BA, Wilkes JG, Du J, Cullen JJ, Buettner GR. 2016. Tumor cells have decreased ability to metabolize H2O2: Implications for pharmacological ascorbate in cancer therapy. Redox Biology 10:274–284. DOI: 10.1016/j.redox.2016.10.010

Doskey CM, Erve TJ van ‘t, Wagner BA, Buettner GR. 2015. Moles of a Substance per Cell Is a Highly Informative Dosing Metric in Cell Culture. PLoS ONE 10. DOI: 10.1371/JOURNAL.PONE.0132572, PMID: 26172833

Du J, Cullen JJ, Buettner GR. 2012. Ascorbic acid: Chemistry, biology and the treatment of cancer. Biochimica et biophysica acta 1826:443. DOI: 10.1016/J.BBCAN.2012.06.003, PMID: 22728050

Du W, Menjivar RE, Donahue KL, Kadiyala P, Velez-Delgado A, Brown KL, Watkoske HR, He X, Carpenter ES, Angeles C V., Zhang Y, Di Magliano MP. 2023. WNT signaling in the tumor microenvironment promotes immunosuppression in murine pancreatic cancer. Journal of Experimental Medicine 220. DOI: /10.1084/JEM.20220503/213546, PMID: 36239683

El Hassouni B, Granchi C, Vallés-Martí A, Supadmanaba IGP, Bononi G, Tuccinardi T, Funel N, Jimenez CR, Peters GJ, Giovannetti E, Minutolo F. 2020. The dichotomous role of the glycolytic metabolism pathway in cancer metastasis: Interplay with the complex tumor microenvironment and novel therapeutic strategies. Seminars in Cancer Biology 60:238–248. DOI: 10.1016/j.semcancer.2019.08.025, PMID: 31445217

Erickson JR, Julie He B, Grumbach IM, Anderson ME. 2011. CaMKII in the cardiovascular system: Sensing redox states. Physiological Reviews 91:889–915. DOI: 10.1152/PHYSREV.00018.2010, PMID: 21742790

Frejno M, Chiozzi RZ, Wilhelm M, Koch H, Zheng R, Klaeger S, Ruprecht B, Meng C, Kramer K, Jarzab A, Heinzlmeir S, Johnstone E, Domingo E, Kerr D, Jesinghaus M, Slotta-Huspenina J, Weichert W, Knapp S, Feller SM, Kuster B. 2017. Pharmacoproteomic characterisation of human colon and rectal cancer. Molecular Systems Biology 13:951. DOI: 10.15252/MSB.20177701, PMID: 29101300

Garrido C, Galluzzi L, Brunet M, Puig PE, Didelot C, Kroemer G. 2006. Mechanisms of cytochrome c release from mitochondria. Cell Death & Differentiation 2006 13:9 13:1423–1433. DOI: 10.1038/sj.cdd.4401950, PMID: 16676004

Ghandi M, Huang FW, Jané-Valbuena J, Kryukov GV, Lo CC, McDonald ER, Barretina J, Gelfand ET, Bielski CM, Li H, Hu K, Andreev-Drakhlin AY, Kim J, Hess JM, Haas BJ, Aguet F, Weir BA, Rothberg MV, Paolella BR, Lawrence MS, Akbani R, Lu Y, Tiv HL, Gokhale PC, De Weck A, Mansour AA, Oh C, Shih J, Hadi K, Rosen Y, Bistline J, Venkatesan K, Reddy A, Sonkin D, Liu M, Lehar J, Korn JM, Porter DA, Jones MD, Golji J, Caponigro G, Taylor JE, Dunning CM, Creech AL, Warren AC, McFarland JM, Zamanighomi M, Kauffmann A, Stransky N, Imielinski M, Maruvka YE, Cherniack AD, Tsherniak A, Vazquez F, Jaffe JD, Lane AA, Weinstock DM, Johannessen CM, Morrissey MP, Stegmeier F, Schlegel R, Hahn WC, Getz G, Mills GB, Boehm JS, Golub TR, Garraway LA, Sellers WR. 2019. Next-generation characterization of the Cancer Cell Line Encyclopedia. Nature 569:503–508. DOI: 10.1038/s41586-019-1186-3

Gillespie M, Jassal B, Stephan R, Milacic M, Rothfels K, Senff-Ribeiro A, Griss J, Sevilla C, Matthews L, Gong C, Deng C, Varusai T, Ragueneau E, Haider Y, May B, Shamovsky V, Weiser J, Brunson T, Sanati N, Beckman L, Shao X, Fabregat A, Sidiropoulos K, Murillo J, Viteri G, Cook J, Shorser S, Bader G, Demir E, Sander C, Haw R, Wu G, Stein L, Hermjakob H, D’Eustachio P. 2022. The reactome pathway knowledgebase 2022. Nucleic Acids Research 50:D687–D692. DOI: 10.1093/NAR/GKAB1028, PMID: 34788843

Goel C, Monga SP, Nejak-Bowen K. 2022. Role and Regulation of Wnt/β-Catenin in Hepatic Perivenous Zonation and Physiological Homeostasis. The American Journal of Pathology 192:4–17. DOI: 10.1016/J.AJPATH.2021.09.007, PMID: 34924168

Günes-Bayir A, Kiziltan HS. 2015. Palliative Vitamin C Application in Patients with Radiotherapy-Resistant Bone Metastases: A Retrospective Study. Nutrition and Cancer. DOI: 10.1080/01635581.2015.1055366, PMID: 26168394

Hänzelmann S, Castelo R, Guinney J. 2013. GSVA: Gene set variation analysis for microarray and RNA-Seq data. BMC Bioinformatics 14. DOI: 10.1186/1471-2105-14-7, PMID: 23323831

Hayes JD, Dinkova-Kostova AT, Tew KD. 2020. Oxidative Stress in Cancer. Cancer Cell 38:167–197. DOI: 10.1016/J.CCELL.2020.06.001, PMID: 32649885

Ho WJ, Jaffee EM, Zheng L. 2020. The tumour microenvironment in pancreatic cancer — clinical challenges and opportunities. Nature Reviews Clinical Oncology 2020 17:9 17:527–540. DOI: 10.1038/s41571-020-0363-5, PMID: 32398706

Huang H, Zhang S, Li Y, Liu Z, Mi L, Cai Y, Wang X, Chen Lishu, Ran H, Xiao D, Li F, Wu J, Li Tingting, Han Q, Chen Liang, Pan X, Li H, Li Tao, He K, Li A, Zhang X, Zhou T, Xia Q, Man J. 2021. Suppression of mitochondrial ROS by prohibitin drives glioblastoma progression and therapeutic resistance. Nature Communications 12. DOI: 10.1038/S41467-021-24108-6, PMID: 34140524

Huang L, Xu AM, Liu W. 2015. Transglutaminase 2 in cancer. American Journal of Cancer Research 5:2756. PMID: 26609482

Kearney AL, Cooke KC, Norris DM, Zadoorian A, Krycer JR, Fazakerley DJ, Burchfield JG, James DE. 2019. Serine 474 phosphorylation is essential for maximal Akt2 kinase activity in adipocytes. The Journal of biological chemistry 294:16729–16739. DOI: 10.1074/JBC.RA119.010036, PMID: 31548312

Keepers YP, Pizao PE, Peters GJ, van Ark-Otte J, Winograd B, Pinedo HM. 1991. Comparison of the sulforhodamine B protein and tetrazolium (MTT) assays for in vitro chemosensitivity testing. European Journal of Cancer and Clinical Oncology 27:897– 900. DOI: 10.1016/0277-5379(91)90142-Z, PMID: 1834124

Kim JH, Hwang S, Lee JH, Im SS, Son J. 2022. Vitamin C Suppresses Pancreatic Carcinogenesis through the Inhibition of Both Glucose Metabolism and Wnt Signaling. International Journal of Molecular Sciences 23:12249. DOI: 10.3390/IJMS232012249/S1, PMID: 36293106

Kim Y, Kim H, Bae S, Choi J, Lim SY, Lee N, Kong JM, Hwang Y, Kang JS, Lee WJ. 2013. Vitamin C Is an Essential Factor on the Anti-viral Immune Responses through the Production of Interferon-α/β at the Initial Stage of Influenza A Virus (H3N2) Infection. Immune Network 13:70. DOI: 10.4110/IN.2013.13.2.70, PMID: 23700397

Kirtonia A, Sethi G, Garg M. 2020. The multifaceted role of reactive oxygen species in tumorigenesis. Cellular and Molecular Life Sciences 77:4459–4483. DOI: 10.1007/S00018-020-03536-5, PMID: 32358622

Kranz RG, Richard-Fogal C, Taylor J-S, Frawley ER. 2009. Cytochrome c Biogenesis: Mechanisms for Covalent Modifications and Trafficking of Heme and for Heme-Iron Redox Control . Microbiology and Molecular Biology Reviews 73:510–528. DOI: 10.1128/MMBR.00001-09/ASSET/017BCAE3-81A9-4F05-984B-377C14A645D8/ASSETS/GRAPHIC/ZMR0030922220007.JPEG, PMID: 19721088

Lane DJR, Richardson DR. 2014. The active role of vitamin C in mammalian iron metabolism: Much more than just enhanced iron absorption! Free Radical Biology and Medicine 75:69–83. DOI: 10.1016/J.FREERADBIOMED.2014.07.007, PMID: 25048971

Lee EE, Ma J, Sacharidou A, Mi W, Salato VK, Nguyen N, Jiang Y, Pascual JM, North PE, Shaul PW, Mettlen M, Wang RC. 2014. A Protein Kinase C Phosphorylation Motif in GLUT1 Affects Glucose Transport and is Mutated in GLUT1 Deficiency Syndrome. Molecular Cell 58:845–853. DOI: 10.1016/j.molcel.2015.04.015, PMID: 25982116

Lee KE, Hahm E, Bae S, Kang JS, Lee WJ. 2017. The enhanced tumor inhibitory effects of gefitinib and L-ascorbic acid combination therapy in non-small cell lung cancer cells. Oncology Letters 14:276–282. DOI: 10.3892/ol.2017.6109, PMID: 28693165

Lee S, Chang J, Blackstone C. 2016. FAM21 directs SNX27–retromer cargoes to the plasma membrane by preventing transport to the Golgi apparatus. Nature Communications 7. DOI: 10.1038/NCOMMS10939, PMID: 26956659

Liberzon A, Subramanian A, Pinchback R, Thorvaldsdóttir H, Tamayo P, Mesirov JP. 2011. Molecular signatures database (MSigDB) 3.0. Bioinformatics 27:1739–1740. DOI: 10.1093/BIOINFORMATICS/BTR260, PMID: 21546393

Linnekamp JF, Van Hooff SR, Prasetyanti PR, Kandimalla R, Buikhuisen JY, Fessler E, Ramesh P, Lee KAST, Bochove GGW, De Jong JH, Cameron K, Van Leersum R, Rodermond HM, Franitza M, Nürnberg P, Mangiapane LR, Wang X, Clevers H, Vermeulen L, Stassi G, Medema JP. 2018. Consensus molecular subtypes of colorectal cancer are recapitulated in in vitro and in vivo models. Cell Death and Differentiation 25:616–633. DOI: 10.1038/s41418-017-0011-5, PMID: 29305587

Liu Y, Huang P, Li Z, Xu C, Wang H, Jia B, Gong A, Xu M. 2022. Vitamin C Sensitizes Pancreatic Cancer Cells to Erastin-Induced Ferroptosis by Activating the AMPK/Nrf2/HMOX1 Pathway. Oxidative Medicine and Cellular Longevity 2022. DOI: 10.1155/2022/5361241, PMID: 35915609

Ma Y, Chapman J, Levine M, Polireddy K, Drisko J, Chen Q. 2014. High-Dose Parenteral Ascorbate Enhanced Chemosensitivity of Ovarian Cancer and Reduced Toxicity of Chemotherapy. Science Translational Medicine 6:222ra18–222ra18. DOI: 10.1126/scitranslmed.3007154, PMID: 24500406

Magrì A, Germano G, Lorenzato A, Lamba S, Chilà R, Montone M, Amodio V, Ceruti T, Sassi F, Arena S, Abrignani S, D’Incalci M, Zucchetti M, Di Nicolantonio F, Bardelli A. 2020. High-dose vitamin C enhances cancer immunotherapy. Science Translational Medicine 12:eaay8707. DOI: 10.1126/scitranslmed.aay8707, PMID: 32102933

Mansoor F, Kumar S, Rai P, Anees F, Kaur N, Devi A, Kumar B, Memon MK, Khan S. 2021. Impact of Intravenous Vitamin C Administration in Reducing Severity of Symptoms in Breast Cancer Patients During Treatment. Cureus 13. DOI: 10.7759/cureus.14867

Michael A, Ainsley A, Joseph A, Jahan N, Michael A, Ainsley A, Joseph A, Jahan N. 2020. First and Second Line Chemotherapeutic Regimens for Non-Small Cell Lung Carcinomas - The Efficacy of Platinum, Non-Platinum and Combination Therapy: A Literature Review. Cureus 12. DOI: 10.7759/CUREUS.11619

Michels AJ, Frei B. 2013. Myths, artifacts, and fatal flaws: Identifying limitations and opportunities in vitamin C research. Nutrients 5:5161–5192. DOI: 10.3390/nu5125161, PMID: 24352093

Miles AL, Burr SP, Grice GL, Nathan JA. 2017. The vacuolar-ATPase complex and assembly factors, TMEM199 and CCDC115, control HIF1α prolyl hydroxylation by regulating cellular iron levels. eLife 6. DOI: 10.7554/ELIFE.22693, PMID: 28296633

Miller MP, Asbury CL, Biggins S. 2016. A TOG Protein Confers Tension Sensitivity to Kinetochore-Microtubule Attachments. Cell 165:1428–1439. DOI: 10.1016/J.CELL.2016.04.030, PMID: 27156448

Mustafi S, Camarena V, Qureshi R, Sant DW, Wilkes Z, Bilbao D, Slingerland J, Kesmodel SB, Wang G. 2021. Vitamin C sensitizes triple negative breast cancer to PI3K inhibition therapy. Theranostics 11:3552. DOI: 10.7150/THNO.53225, PMID: 33664847

Ngo B, Van Riper JM, Cantley LC, Yun J. 2019. Targeting cancer vulnerabilities with high-dose vitamin C. Nature Reviews Cancer 1. DOI: 10.1038/s41568-019-0135-7

Oliveros, J.C. 2015. Venny. An interactive tool for comparing lists with Venn’s diagrams.

Perez-Riverol Y, Bai J, Bandla C, García-Seisdedos D, Hewapathirana S, Kamatchinathan S, Kundu DJ, Prakash A, Frericks-Zipper A, Eisenacher M, Walzer M, Wang S, Brazma A, Vizcaíno JA. 2022. The PRIDE database resources in 2022: a hub for mass spectrometry-based proteomics evidences. Nucleic Acids Research 50:D543. DOI: 10.1093/NAR/GKAB1038, PMID: 34723319

Piersma SR, Fiedler U, Span S, Lingnau A, Pham T V., Hoffmann S, Kubbutat MHG, Jiménez CR. 2010. Workflow comparison for label-free, quantitative secretome proteomics for cancer biomarker discovery: Method evaluation, differential analysis, and verification in serum. Journal of Proteome Research 9:1913–1922. DOI: 10.1021/PR901072H/SUPPL_FILE/PR901072H_SI_008.PDF, PMID: 20085282

Pin G, Huanting L, Chengzhan Z, Xinjuan K, Yugong F, Wei L, Shifang L, Zhaojian L, Kun H, Weicheng Y, Yingying L, Yongming Q, Yanan Y. 2020. Down-Regulation of PDCD4 Promotes Proliferation, Angiogenesis and Tumorigenesis in Glioma Cells. Frontiers in Cell and Developmental Biology 8:593685. DOI: 10.3389/FCELL.2020.593685/BIBTEX

Pizzo V, Lendeckel U, Karimi F, Al Abdulla R, Wolke C. 2023. The Role of the Ectopeptidase APN/CD13 in Cancer. Biomedicines 2023, Vol. 11, Page 724 11:724. DOI: 10.3390/BIOMEDICINES11030724

Polireddy K, Dong R, Reed G, Yu J, Chen P, Williamson S, Violet P-C, Pessetto Z, Godwin AK, Fan F, Levine M, Drisko JA, Chen Q. 2017. High Dose Parenteral Ascorbate Inhibited Pancreatic Cancer Growth and Metastasis: Mechanisms and a Phase I/IIa study. Scientific Reports 7:17188. DOI: 10.1038/s41598-017-17568-8

Porporato PE, Filigheddu N, Pedro JMBS, Kroemer G, Galluzzi L. 2017. Mitochondrial metabolism and cancer. Cell Research 2017 28:3 28:265–280. DOI: 10.1038/CR.2017.155, PMID: 29219147

Qin S, Wang G, Chen L, Geng H, Zheng Y, Xia C, Wu S, Yao J, Deng L. 2023. Pharmacological vitamin C inhibits mTOR signaling and tumor growth by degrading Rictor and inducing HMOX1 expression. PLOS Genetics 19:e1010629. DOI: 10.1371/JOURNAL.PGEN.1010629, PMID: 36787291

Raudvere U, Kolberg L, Kuzmin I, Arak T, Adler P, Peterson H, Vilo J. 2019. g:Profiler: a web server for functional enrichment analysis and conversions of gene lists (2019 update). Nucleic Acids Research 47:W191–W198. DOI: 10.1093/NAR/GKZ369, PMID: 31066453

Reczek CR, Chandel NS. 2015. Revisiting vitamin C and cancer. Science 350:1317–1318. DOI: 10.1126/science.aad8671, PMID: 26659042

Schoenfeld JD, Sibenaller ZA, Mapuskar KA, Wagner BA, Cramer-Morales KL, Furqan M, Sandhu S, Carlisle TL, Smith MC, Abu Hejleh T, Berg DJ, Zhang J, Keech J, Parekh KR, Bhatia S, Monga V, Bodeker KL, Ahmann L, Vollstedt S, Brown H, Shanahan Kauffman EP, Schall ME, Hohl RJ, Clamon GH, Greenlee JD, Howard MA, Shultz MK, Smith BJ, Riley DP, Domann FE, Cullen JJ, Buettner GR, Buatti JM, Spitz DR, Allen BG. 2017. O2⋅− and H2O2-Mediated Disruption of Fe Metabolism Causes the Differential Susceptibility of NSCLC and GBM Cancer Cells to Pharmacological Ascorbate. Cancer Cell 31:487–500.e8. DOI: 10.1016/J.CCELL.2017.02.018, PMID: 28366679

Sergushichev AA. 2016. An algorithm for fast preranked gene set enrichment analysis using cumulative statistic calculation. bioRxiv 060012. DOI: 10.1101/060012

Sinha BK, van ‘t Erve TJ, Kumar A, Bortner CD, Motten AG, Mason RP. 2017. Synergistic enhancement of topotecan-induced cell death by ascorbic acid in human breast MCF-7 tumor cells. Free Radical Biology and Medicine. DOI: 10.1016/j.freeradbiomed.2017.10.377, PMID: 29079526

Sorenson GD, Pettengill S, Brinck-Johnsen T, Cate CC, Maurer LH. 1981. Hormone Production by Cultures of Small-Cell Carcinoma of the Lung. Cancer. DOI: 10.1002/1097-0142

Soula M, Weber RA, Zilka O, Alwaseem H, La K, Yen F, Molina H, Garcia-Bermudez J, Pratt DA, Birsoy K. 2020. Metabolic determinants of cancer cell sensitivity to canonical ferroptosis inducers. Nature Chemical Biology 2020 16:12 16:1351–1360. DOI: 10.1038/s41589-020-0613-y, PMID: 32778843

Su X, Li P, Han B, Jia H, Liang Q, Wang H, Gu M, Cai J, Li S, Zhou Y, Yi X, Wei W. 2021. Vitamin C sensitizes BRAFV600E thyroid cancer to PLX4032 via inhibiting the feedback activation of MAPK/ERK signal by PLX4032. Journal of Experimental and Clinical Cancer Research 40. DOI: 10.1186/s13046-021-01831-y, PMID: 33468157

Su X, Shen Z, Yang Q, Sui F, Pu J, Ma J, Ma S, Yao D, Ji M, Hou P. 2019. Vitamin C kills thyroid cancer cells through ROS-dependent inhibition of MAPK/ERK and PI3K/AKT pathways via distinct mechanisms. Theranostics. DOI: 10.7150/thno.35219, PMID: 31285773

Subramanian A, Tamayo P, Mootha VK, Mukherjee S, Ebert BL, Gillette MA, Paulovich A, Pomeroy SL, Golub TR, Lander ES, Mesirov JP. 2005. Gene set enrichment analysis: A knowledge-based approach for interpreting genome-wide expression profiles. Proceedings of the National Academy of Sciences of the United States of America 102:15545–15550. DOI: 10.1073/PNAS.0506580102/SUPPL_FILE/06580FIG7.JPG, PMID: 16199517

Sullivan LB, Chandel NS. 2014. Mitochondrial reactive oxygen species and cancer. Cancer and Metabolism 2. DOI: 10.1186/2049-3002-2-17, PMID: 25671107

Supek F, Bošnjak M, Škunca N, Šmuc T. 2011. REVIGO Summarizes and Visualizes Long Lists of Gene Ontology Terms. PLoS ONE 6:e21800. DOI: 10.1371/journal.pone.0021800

Tanner JJ, Fendt SM, Becker DF. 2018. The Proline Cycle As a Potential Cancer Therapy Target. Biochemistry 57:3433–3444. DOI: 10.1021/ACS.BIOCHEM.8B00215/ASSET/IMAGES/MEDIUM/BI-2018-00215E_0006.GIF, PMID: 29648801

Vallés-Martí A, Mantini G, Manoukian P, Waasdorp C, Sarasqueta AF, Haas RR de G, Henneman AA, Piersma SR, Pham T V., Knol JC, Giovannetti E, Bijlsma MF, Jiménez CR. 2023. Phosphoproteomics guides effective low-dose drug combinations against pancreatic ductal adenocarcinoma. Cell Reports 42:112581. DOI: 10.1016/J.CELREP.2023.112581, PMID: 37269289

van der Reest J, Gottlieb E. 2016. Anti-cancer effects of vitamin C revisited. Cell Research 2016 26:3 26:269–270. DOI: 10.1038/cr.2016.7

Vera JC, Rivas CI, Fischbarg J, Golde DW. 1993. Mammalian facilitative hexose transporters mediate the transport of dehydroascorbic acid. Nature 364:79–82. DOI: 10.1038/364079a0

Vichai V, Kirtikara K. 2006. Sulforhodamine B colorimetric assay for cytotoxicity screening. Nature Protocols 2006 1:3 1:1112–1116. DOI: 10.1038/nprot.2006.179, PMID: 17406391

Wang L, Luo X, Li C, Huang Y, Xu P, Lloyd-Davies LH, Delplancke T, Peng C, Gao R, Qi H, Tong C, Baker P. 2017. Triethylenetetramine Synergizes with Pharmacologic Ascorbic Acid in Hydrogen Peroxide Mediated Selective Toxicity to Breast Cancer Cell. Oxidative Medicine and Cellular Longevity. DOI: 10.1155/2017/3481710, PMID: 28280522

Wang X, Xu S, Zhang L, Cheng X, Yu H, Bao J, Lu R. 2021. Vitamin C induces ferroptosis in anaplastic thyroid cancer cells by ferritinophagy activation. Biochemical and biophysical research communications 551:46–53. DOI: 10.1016/J.BBRC.2021.02.126, PMID: 33714759

Welsh JL, Wagner BA, van’t Erve TJ, Zehr PS, Berg DJ, Halfdanarson TR, Yee NS, Bodeker KL, Du J, Roberts LJ, Drisko J, Levine M, Buettner GR, Cullen JJ. 2013. Pharmacological ascorbate with gemcitabine for the control of metastatic and node-positive pancreatic cancer (PACMAN): results from a phase I clinical trial. Cancer Chemotherapy and Pharmacology 71:765–775. DOI: 10.1007/s00280-013-2070-8, PMID: 23381814

Xu J, Jia Z, Knutson MD, Leeuwenburgh C. 2012. Impaired Iron Status in Aging Research. International Journal of Molecular Sciences 2012, Vol. 13, Pages 2368-2386 13:2368–2386. DOI: 10.3390/IJMS13022368, PMID: 22408459

Yang G, Yan Y, Ma Y, Yang Y. 2017. Vitamin C at high concentrations induces cytotoxicity in malignant melanoma but promotes tumor growth at low concentrations. Molecular Carcinogenesis. DOI: 10.1002/mc.22654, PMID: 28370562

Yun J, Mullarky E, Lu C, Bosch KN, Kavalier A, Rivera K, Roper J, Chio IIC, Giannopoulou EG, Rago C, Muley A, Asara JM, Paik J, Elemento O, Chen Z, Pappin DJ, Dow LE, Papadopoulos N, Gross SS, Cantley LC. 2015. Vitamin C selectively kills KRAS and BRAF mutant colorectal cancer cells by targeting GAPDH. Science 350:1391–1396. DOI: 10.1126/science.aaa5004

Zeng LH, Wang QM, Feng LY, Ke YD, Xu QZ, Wei AY, Zhang C, Ying RB. 2019. High-dose vitamin C suppresses the invasion and metastasis of breast cancer cells via inhibiting epithelial-mesenchymal transition. OncoTargets and Therapy 12:7405–7413. DOI: 10.2147/OTT.S222702

Zhao L, Quan Y, Wang J, Wang F, Zheng Y, Zhou A. 2015. Vitamin C inhibit the proliferation, migration and epithelial-mesenchymal-transition of lens epithelial cells by destabilizing HIF-1α. International Journal of Clinical and Experimental Medicine. PMID: 26628999

Zwolak I, Wnuk E, Świeca M. 2022. Identification of Potential Artefacts in In Vitro Measurement of Vanadium-Induced Reactive Oxygen Species (ROS) Production. International Journal of Environmental Research and Public Health 19:15214. DOI: 10.3390/ijerph192215214

